# Effects of Familial Alzheimer’s Disease Mutations on the Folding Free Energy and Dipole-Dipole Interactions of the Amyloid *β*-Peptide

**DOI:** 10.1101/2022.05.19.492750

**Authors:** Darcy S. Davidson, Joshua A. Kraus, Julia M. Montgomery, Justin A. Lemkul

## Abstract

Familial Alzheimer’s disease (FAD) mutations of the amyloid *β*-peptide (A*β*) are known to lead to early onset and more aggressive Alzheimer’s disease. FAD mutations such as “Iowa” (D23N), “Arctic” (E22G), “Italian” (E22K), and “Dutch” (E22Q) have been shown to accelerate A*β* aggregation relative to the wild-type (WT). The mechanism by which these mutations facilitate increased aggregation is unknown, but each mutation results in a change in net charge of the peptide. Previous studies have used nonpolarizable force fields to study A*β*, providing some insight into how this protein unfolds. However, nonpolarizable force fields have fixed charges that lack the ability to redistribute in response to changes in local electric fields. Here, we performed polarizable molecular dynamics (MD) simulations on the full-length A*β*_42_ of WT and FAD mutations and calculated folding free energies of the A*β*_15-27_ fragment via umbrella sampling. By studying both the full-length A*β*_42_ and a fragment containing mutations and the central hydrophobic cluster (residues 17-21), we were able to systematically study how these FAD mutations impact secondary and tertiary structure and the thermodynamics of folding. Electrostatic interactions, including those between permanent and induced dipoles, affected sidechain properties, salt bridges, and solvent interactions. The FAD mutations resulted in shifts in the electronic structure and solvent accessibility at the central hydrophobic cluster and the hydrophobic C-terminal region. Using umbrella sampling, we found that the folding of the WT and E22 mutants are enthalpically driven, whereas the D23N mutant is entropically driven, arising from a different unfolding pathway and peptide-bond dipole response. Together, the unbiased, full-length and umbrella sampling simulations of fragments reveal that the FAD mutations perturb nearby residues and others in hydrophobic regions to potentially alter solubility. These results highlight the role electronic polarizability plays in amyloid misfolding and the role of heterogeneous microenvironments that arise as conformational change takes place.

## Introduction

Alzheimer’s Disease (AD) is a neurodegenerative disease that causes irreversible and progressive brain damage.^1^ There are two proteins believed to be central to the cytotoxicity in AD, the microtubule-associated protein tau and the amyloid *β*-peptide (A*β*), though the exact mechanism(s) by which these proteins contribute to disease are not fully understood. ^2^ It is known that A*β* unfolds, aggregates, then self-assembles into highly ordered insoluble extracellular A*β* senile plaques. A*β* is derived from the amyloid precursor protein (APP), a membrane protein believed to be associated with neural plasticity.^3^ APP is cleaved by *β*-secretase and *γ*-secretase to form A*β*, along with a soluble extracellular protein and a C-terminal fragment.^4,5^ The *γ*-secretase cleavage is processive and can be initiated at several positions in the C-terminal fragment, resulting in A*β* polypeptides containing 36-43 amino acids.^4,5^ The more extended forms of A*β* are more hydrophobic and fibrillogenic; specifi-cally, the 42-residue alloform of A*β* is more abundant in the brains of AD patients. ^6,7^ After *γ*-secretase cleavage, A*β* is released into the extracellular space where it aggregates into low-molecular-weight oligomers and ultimately fibrils and plaques.

Amyloid fibrils are thought to be an end state of the A*β* aggregation cascade that contributes to the pathogenesis of AD. ^8^ Thus, studying A*β* as a monomer and in oligomers can provide us with valuable insight into the earliest events in AD, which may lead to new therapeutic approaches. The A*β* sequence is characterized by alternating regions of polarity and hydrophobicity. Previous work has proposed that the hydrophilic N-terminal region of A*β* shields the hydrophobic C-terminal region from water, thereby promoting fibrillogenesis.^9^ In addition, residues 17-21, known as the “central hydrophobic cluster” (CHC), are also thought to be a nucleation site for A*β* aggregation.^10^ The CHC is flanked by polar residues, H_13_HQK_16_ on the N-terminal side, and the acidic dyad (residues E22 and D23) on the C-terminal side, which may promote the formation of a network of specific interactions that enhance aggregation. Ultimately, the atomistic details of A*β* unfolding and aggregation are poorly understood, and it is unclear how low-molecular-weight oligomers form and lead to fibril structures.

Several genetic mutations occur in A*β*, including the D23N “Iowa,”^11,12^ E22G “Arctic,”^13,14^ E22K “Italian,”^15^ and E22Q “Dutch”^16–18^ variants that lead to familial AD (FAD), forms of AD in which symptoms start as early as 35 years of age. ^11,12^ These mutations are known to increase the severity of disease, and on the molecular level, enhance the kinetics of A*β* aggregation.^11,14,15,19^ These mutations are also known to increase the rate of secondary nucleation due to the changes in charge and electrostatic nature of the peptide. ^19^ The mechanism by which mutations facilitate increased aggregation and a more severe form of AD is unknown. One recent simulation study applied a coarse-grain model to determine that A*β*_42_ monomers are more prone to adopting fibril-like structures, and that FAD mutations in A*β*_40_ monomers reduce the free energy barriers between monomer and fibril-like structures. ^20^ Thus, these mutations may play a role in modulating A*β* structure on the monomer level.

A*β* is difficult to investigate experimentally due to its propensity to aggregate in solution.^21^ Thus, many theoretical investigations have been carried out on this peptide using molecular dynamics (MD) simulations. Many of these studies, as well as corresponding experimental investigations for A*β* and a wide range of amyloidogenic proteins, have recently been reviewed.^22^ Extensive effort has been devoted to simulating A*β*, including full-length and fragment simulations of monomers and aggregates of varying sizes and compositions. It is impossible to summarize all of these efforts comprehensively in this space, but several recent findings from MD simulations on the A*β* monomer are important to note here. An early simulation study of the A*β*_10-35_ fragment found good agreement with NMR data that suggested an important bend structure.^23^ Though these simulations were very short (1 ns) by current standards, they provided a foundation for subsequent work to interrogate the agreement of simulation and experimental outcomes. A more recent simulation study by Granata et al. found that A*β*_40_ is fully disordered in its lowest energy conformers, with residual secondary structure only populated in higher-energy states.^24^

Considerable effort has gone into studying the aggregation pathway and kinetics of A*β* and the FAD mutants.^21,25^ Previous MD simulations have provided insights into the atomistic details of amyloid dynamics, and nearly all of these studies have used nonpolarizable force fields (FFs) to study A*β*, providing some details as to how these proteins unfold. ^26,27^ However, some discrepancies have been noted, mainly due to inadequate sampling or shortcomings in the FFs.^27–29^ One potential issue is that nonpolarizable FFs have fixed charges that lack the ability to redistribute in response to changes in their environment. As such, nonpolarizable FFs may face challenges in modeling complex microenvironments and balance intra-protein and protein-water interactions. We note, however, the success of recently developed FFs such as a99SB-*disp* that has been specifically optimized to model intrinsically disordered proteins.^30^ While a99SB-*disp* was stated to perform well in simulations of A*β*_40_, Paul et al. recently concluded that a different AMBER force field variant, a99SB-UCB, was the most suitable for modeling A*β*_40_,^31^ based on its ability to better model kinetics of structural interconversion, chemical shifts, and FRET-derived structural quantities. The a99SB-UCB force field is a modification of the AMBER ff99SB FF^32^ for use with the TIP4P-Ew water model,^33^ and includes refinements of backbone dihedral parameters^34^ and protein-water Lennard-Jones terms.^35^ These FF terms are often that targets of optimization, in response to the inherent difficulty in simultaneously balancing conformational sampling and appropriate strength of intra-protein and protein-solvent interactions. It is for this reason why it is beneficial to extend simulation efforts to include polarizable FFs, which may be more sensitive to these behaviors.

We previously performed simulations of an A*β* fibril to understand the factors that stabilize tertiary and quaternary interactions.^36^ We found that amino acid sidechain dipole moments were responsive to their local environments and that unusual microenvironments and confined water molecules may play a role in fibril stabilization. The extent to which these properties extend to A*β* monomers is unknown, but the importance of electronic polarization and charged amino acids has become apparent. Moreover, in 2015, we showed that induced polarization effects among FAD mutations in the A*β*_15-27_ fragment modulate unfolding of this peptide, behaviors that were explicitly linked to properties of different amino-acid sidechains.^37^ Based on these previous studies, it is important to expand our understanding of polarization effects in A*β* to include full-length monomers to investigate conformational sampling and to quantify differences in folding free energies of critical structural motifs. To this end, we performed polarizable MD simulations of the 42-residue A*β* monomer (A*β*_42_), including wild-type (WT), D23N, E22G, E22K, and E22Q variants. We further applied umbrella sampling (US) to investigate the helix-coil transition of the fragment encompassing residues 15-27 of A*β* investigated previously^37^ to determine if mutations directly impact the ability of this fragment to adopt a stable *α*-helix. By studying both the fragment and fulllength peptides, we sought to contextualize our findings and gain further insight into how charge modulates A*β* unfolding and potentially how these mutations alter the aggregation pathway by shifting the properties of the monomers.

## Methods

### Full-length A*β*_42_ system construction

The starting coordinates for the WT A*β*_42_ were obtained from the first model of the NMR ensemble deposited in PDB 1IYT.^38^ The mutations (E22G, E22K, E22Q, and D23N) were introduced by deleting the relevant sidechain atoms and rebuilding the new mutated side chain using the CHARMM internal coordinate builder.^39^ The termini of all WT and mutant peptides were modeled in their ionized forms. We simulated 3 replicates of the WT and mutants for 1 *μ*s each, totaling 3 *μ*s for each A*β*_42_ variant. As will be described below, these simulations led to poor agreement with experimental properties of the peptide, motivating the generation of non-helical starting states.

### System setup and general molecular dynamics protocol

All systems were initially prepared using the additive CHARMM36m^29^ (C36m) protein FF, then later converted to the Drude-2019 FF.^40^ Each protein was solvated in a cubic box of CHARMM-modified TIP3P^41–43^ water and ∼150 mM KCl, including K^+^ counterions (Table 1). The systems were energy-minimized in CHARMM^39^ using 1000 steps of steepest descent minimization and 1000 steps of adopted-basis Newton-Raphson (ABNR) minimization. Following minimization, equilibration was carried out in NAMD^44^ for 1 ns. During equilibration, position restraints were applied to all non-hydrogen protein atoms with a force constant of 5 kcal mol^-1^ Å^-2^. An NPT ensemble was maintained by regulating temperature at 310 K with the Langevin thermostat method^45,46^ using a friction coefficient, *γ*, of 5 ps^-1^. The pressure was set to 1 atm using the Langevin piston method^46^ (oscillation period = 200 fs; decay time = 100 fs). Periodic boundary conditions were applied in all directions, and the short-range van der Waals forces were smoothly switched to zero over 10 – 12 Å. The particle mesh Ewald (PME) method^47,48^ was used to calculate the electrostatic interactions with a real-space cutoff of 12 Å and grid spacing of approximately 1 Å. Bonds to hydrogen atoms were held rigid using the SHAKE^49^ algorithm, allowing an integration time step of 2 fs in C36m simulations. Three independent simulations were performed for each peptide system by generating different starting velocities at the beginning of equilibration.

**Table 1:**
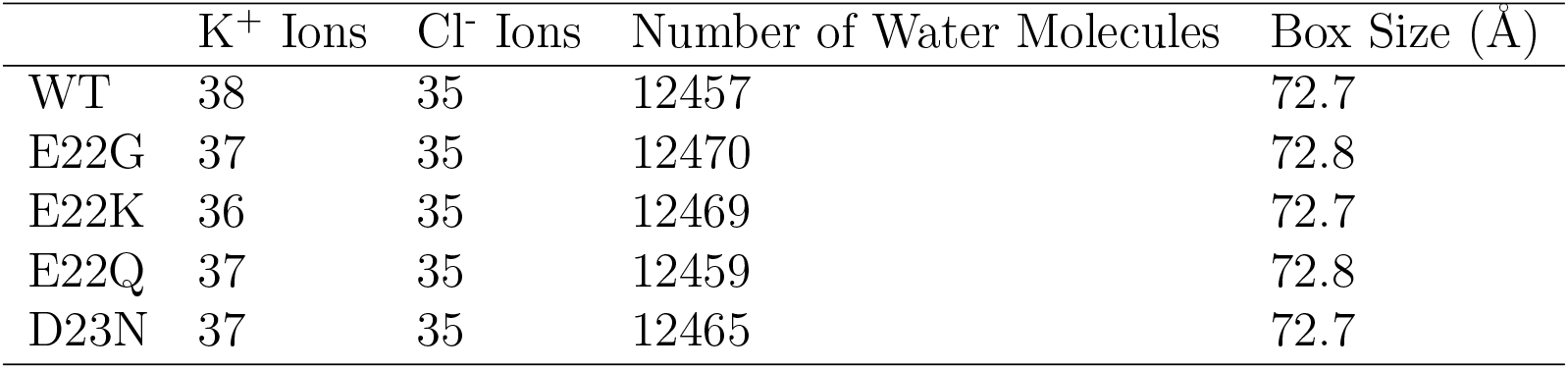
Contents of full-length A*β*_42_ simulation systems. The box size reflects the edge length of the unit cell after NPT equilibration.

Following C36m equilibration, the systems were converted to the Drude polarizable model using CHARMM by adding Drude oscillators and lone pairs to the equilibrated coordinates. Additionally, the TIP3P water molecules were converted to the polarizable SWM4-NDP^50^ model. Drude oscillators were relaxed using 2000 steps of steepest descent and 1000 steps of ABNR energy minimization. The Drude systems were equilibrated in NAMD using extended Lagrangian integration, implemented as Langevin dynamics.^51,52^ An NPT ensemble was maintained, and the temperature and pressure were set to 310 K and 1 atm, respectively, unless otherwise specified. Bonds involving hydrogen atoms were constrained using the SHAKE algorithm as described above. Temperature was regulated using a dual Langevin thermostat, coupling all real atoms at 310 K (*γ* = 5 ps^-1^) and the Drude oscillators to a low-temperature relative thermostat at 1 K (*γ* = 20 ps^-1^). The short-range Lennard-Jones potential was switched to zero from 10 – 12 Å and electrostatic interactions were calculated with PME, using the same settings as the additive simulations. The same harmonic position restraints and bond constraints were applied as the additive system, but the integration was set to 1 fs because of the high-frequency nature of the Drude-atom bonds. To avoid polarization catastrophe, a “hard wall” constraint^53^ was applied to allow a maximum Drude-atom bond length of 0.2 Å. Equilibration of polarizable systems was carried out for 1 ns. Following equilibration, the restraints were removed and the production simulations were performed in OpenMM.^54,55^ The NPT ensemble was maintained using the same thermostat settings as equilibration, except that pressure was regulated at 1 atm using a Monte Carlo barostat with box scaling attempted every 25 integration steps.

### Full-length, melted A*β*_42_ system construction

To generate disordered starting states for full-length A*β*_42_, we simulated the protein using the C36m FF in triplicate at 500 K. Following energy minimization, the systems were equilibrated in NAMD using an NVT ensemble to prevent spurious solvent expansion that may occur with an NPT ensemble at elevated temperature. Equilibration was performed as described in the general methods section, except that the reference temperature for the Langevin piston thermostat was set to 500 K. Unrestrained simulations were performed in OpenMM, using the Andersen method to regulate temperature with a collision frequency of 1 ps^-1^. Unrestrained simulations were performed for 100 ns (WT, E22G, E22K, E22Q, and D23N) or 200 ns in one of the three E22Q replicate system, starting from helical conformers as described above (“Full-length A*β* system construction”). We simulated the E22Q peptide for a longer time because the peptide bond between His6 and Asp7 isomerized into the *cis* conformation in the first 100 ns of the simulation. Thus, we extended this simulation for an additional 100 ns to allow the peptide bond to revert back into the *trans* configuration. We calculated the backbone RMSD for each replicate and extracted the snapshot with the highest value, thus generating three distinct conformations for each system. These denatured structures were then converted to the Drude model as described above and simulated at 310 K in triplicate for 1 *μ*s each, totaling 3 *μ*s for each system.

### Umbrella sampling protocol

The initial coordinates for US simulations of the A*β* fragment 15-27 (A*β*_15-27_) were taken from previous work.^37^ To negate the end effects from charged termini, we capped the N-and C-termini of the A*β*_15-27_ fragment with acetyl and N-methylamide groups, respectively. The WT and mutant peptide fragment systems were prepared as described in the general methods section, and were initially prepared with the additive model and solvated in a 65-Å cubic box (Table 2). To calculate ΔH and ΔS using the van’t Hoff equation, we performed each series of US simulations for each peptide at 298 K, 304 K, 310 K, 316 K, and 350 K with Drude-2019.

**Table 2:** Contents of A*β*_15-27_ US simulation systems. The box size reflects the edge length of the unit cell after NPT equilibration.

|  | K<sup>+</sup> Ions | Cl<sup>-</sup> Ions | Number of Water Molecules | Box Size (Å) |
| --- | --- | --- | --- | --- |
| WT<sub>15-27</sub> | 25 | 24 | 8534 | 63.7 |
| E22G<sub>15-27</sub> | 25 | 25 | 8523 | 63.7 |
| E22K<sub>15-27</sub> | 24 | 25 | 8517 | 63.8 |
| E22Q<sub>15-27</sub> | 25 | 25 | 8515 | 63.7 |
| D23N<sub>15-27</sub> | 25 | 25 | 8517 | 63.7 |

Following equilibration, systems were then prepared for US by creating 32 windows along the reaction coordinate, defined as the distance between the C*α* atoms of Gln15 and Asn27. Each window increased the distance along the reaction coordinate by 0.5 Å, starting at 18 Å ending with 33.5 Å. The spring constant, *k*, was set to 10 kcal mol^-1^ Å^-2^. US was carried out in OpenMM and the windows were simulated for 100 ns (WT and E22K) or 150 ns (E22G, E22K, and E22Q) such that all systems reached convergence, which we defined as obtaining ΔG_fold_ that was statistically invariant over a contiguous 50-ns block of time. We computed ΔG_fold_ in 5-ns intervals and plotted these values over time to compute the average and standard deviation over 50 ns. Thus, if each of the 5-ns interval values of ΔG_fold_ fell within the standard deviation about the average, we concluded the simulations were converged. If any value of ΔG_fold_ for a given 5-ns interval fell outside this range, but the mean did not shift outside the existing range of values, we also considered the simulations converged.

### Structural and C*α* chemical shift analysis

We compared our full-length and melted WT structures to experimental data by calculating the secondary structure, radius of gyration (R_g_), and C*α* chemical shifts. Secondary structure was calculated according to the DSSP method,^56^ as implemented in GROMACS. We used standard CHARMM analysis tools to calculate R_g_. Chemical shifts were calculated by using the SPARTA+ software, ^57^ analyzing snapshots every 100 ps, and then calculating an ensemble average. The calculated C*α* chemical shifts were then compared against the C*α* chemical shifts reported by Hou et al.^58^

### Clustering analysis

To better characterize the subpopulations of conformers in the simulations of full-length A*β*_42_, we performed 2D-clustering using the clustering function in CHARMM.^59–61^ Replicate simulations were pooled and snapshots were separated into groups based on a self-defined maximum radius of 3 Å. The two dimensions used for clustering were the backbone RMSD and the protein R_g_.

### US analysis methods

Following the US simulations, we calculated the free energy by using the WHAM algorithm.^62,63^ We set the bin size to 0.05 Å and a tolerance of 0.001 kcal mol^-1^ Å^-1^. Calculation of the folding free energy (ΔG_fold_) requires a distinction to be made between folded and unfolded states. To do so, we calculated the percentage of *α*-helicity for each window, setting a cutoff value of 50% to assign folded (above the cutoff) and unfolded (below the cutoff). Thus, multiple windows contribute to the unfolded and folded ensembles. ΔG_fold_ was then computed according to Eq. 1:

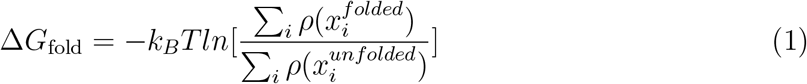

To calculate the individual contributions of ΔH and ΔS to ΔG_fold_, we performed simulations at various temperatures, then calculated the ΔG_fold_ for each. From this quantity, we computed the equilibrium constant, K, which was plotted as a function of inverse temperature according to the van’t Hoff relation (Eq. 2):

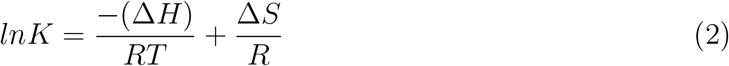

As will be discussed below, we applied the linear van’t Hoff relation to the WT, E22G, E22K, and E22Q systems. For the D23N mutant, we applied the quadratic form of the van’t Hoff^64^ relation (Eq. 3):

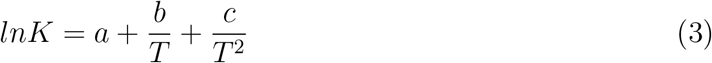

The values of ΔH and ΔS can thus be calculated according to Eqs. 4 and 5:

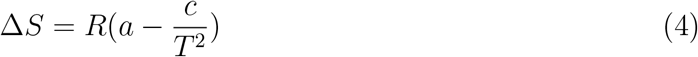

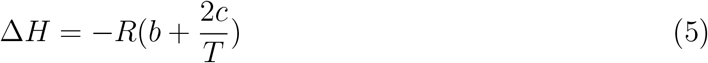

### Dipole moment analysis

Dipole moments of various components of protein structures were calculated using built-in CHARMM functions. Peptide-bond dipole enhancement analysis was performed as described by Huang and MacKerell.^65^

## Results and Discussion

The effects of the FAD mutations on the conformational ensemble of monomeric A*β*_42_ are not fully understood. Changes in microenvironment and charge are thought to play a key role in how proteins fold. ^36,66^ Since the FAD mutations studied here are charge-altering mutations, we expected that using a polarizable FF would provide valuable insights into how changes in charge alter the conformational ensemble of A*β*_42_. We simulated both the full-length peptide and a key fragment to gain a greater understanding of the role that charge plays on dictating secondary and tertiary structure, and the extent to which the mutations alter the free energy difference between helical and disordered states. Studying the fulllength peptide is important because prior work suggests that the polar N-terminal residues (particularly residues 1-10) are necessary for cytotoxicity and aggregation.^67^ In addition, understanding the tertiary structure and contacts that are formed provides insight into how A*β*_42_ initially unfolds and how mutations modulate exposure of individual residues to water and thus impact the aggregation pathway.

### WT A*β*_42_ structural validation

We initially performed simulations of the WT A*β*_42_ peptide starting from the mostly *α*-helical configuration deposited in PDB entry 1IYT.^38^ To assess the validity of the simulated ensemble, we calculated the secondary structure content, R_g_, and C*α* chemical shifts and compared these results to experimental values. The calculated *α*-helicity was 72 ± 15%, which is substantially higher than the experimentally determined value of 5%.^68^ The sim-ulated R_g_ was 1.6 ± 0.2 nm, higher than one experimental value of 0.9 ± 0.1 nm,^69^ but consistent with a more recent measurement of 1.59 ± 0.01 nm from FRET experiments.^70^

We then compared the C*α* chemical shifts computed with SPARTA+ to the values reported by Hou et al.,^58^ finding that values were systematically up-shifted (a reflection of the aberrantly large *α*-helicity, Figure 1A). The RMSD of the simulated C*α* shifts relative to experimental values was 2.5 ppm, with an R^2^ value of 0.96. Together, these analyses suggested the peptide remained far too helical even if the R_g_ value was in line with recent experimental data. The 1IYT structure was determined in a solvent mixture that included hexafluoroisopropanol,^38^ leading to a structure that is higher in *α*-helicity than what is observed for A*β*_42_ in water.^68^ One challenge in performing polarizable MD simulations is that the SWM4-NDP water model diffuses more slowly than additive models like TIP3P, but is more accurate in this regard.^50^ Thus, observing the loss of the helical structure of the peptide may not be feasible in unbiased simulations.

**Figure 1:**
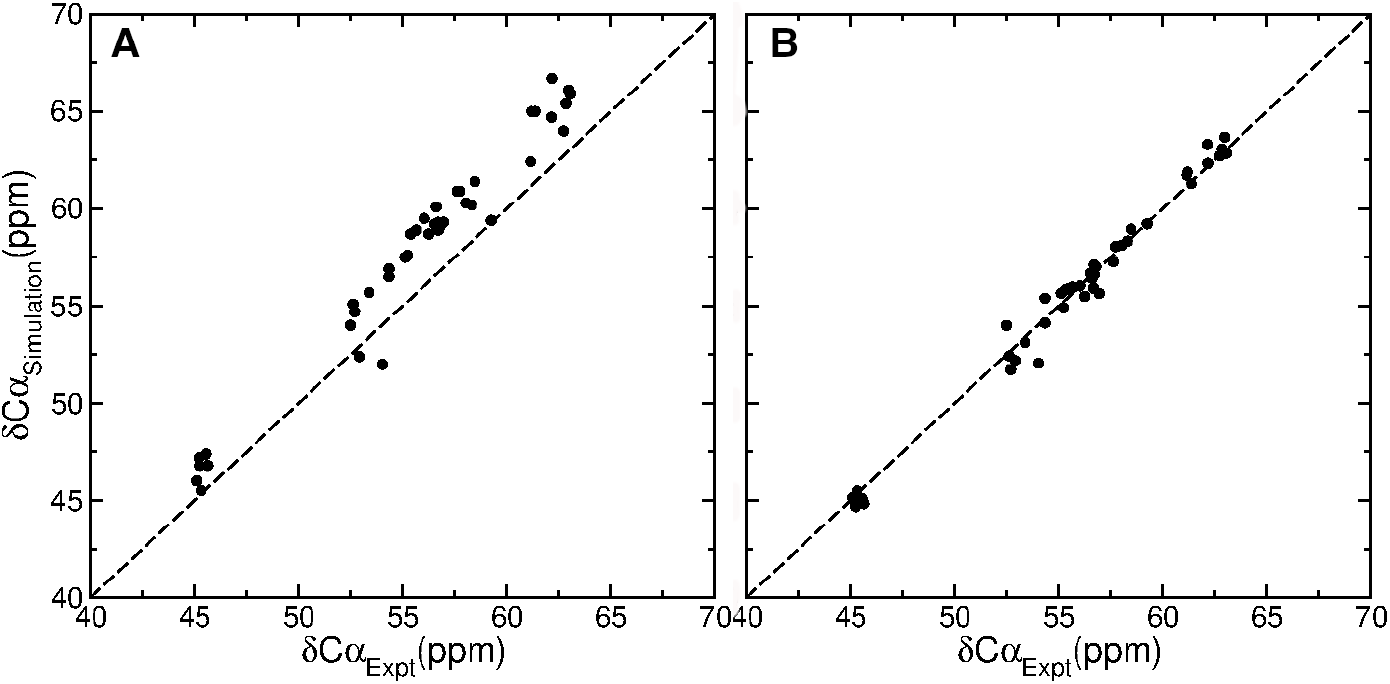
Chemical shift comparison of simulations of the WT A*β*_42_ peptide initiated from (A) the 1IYT structure and (B) heat-denatured configurations.

To circumvent this problem, we denatured the initially helical A*β*_42_ peptide from PDB 1IYT by performing simulations with the C36m force field at 500 K for 100-200 ns. Starting from disordered states, we performed new simulations at 310 K and compared the resulting conformational ensemble of the WT A*β*_42_ peptide to available experimental data. The *α*-helical content of the disordered ensemble was 5 ± 5%, which is consistent with experimental results. The R_g_ was 1.2 ± 0.2 nm, which is slightly higher than the value obtained by Nag et al.^69^ but lower than that determined by Meng et al.^70^ Importantly, there is still some disagreement about the actual R_g_ for A*β*_42_, as Meng et al. note that prior estimates of R_g_ of 1.0-1.2 nm, often obtained in simulations, are consistent with a wide range of NMR data (chemical shifts, J-couplings, and NOE distances). Therefore, some ambiguity remains on what value of R_g_ should be used as a standard for assessing these simulation results. Nevertheless, the R_g_ value obtained in our simulations is in line with previous simulation studies and is within a range defined by multiple experiments.^69,70^

The calculated C*α* chemical shifts were more consistent in this disordered ensemble than those produced by simulations starting from the 1IYT structure, with an RMSD of only 0.6 ppm and R^2^ = 0.99 (Figure 1B). Thus, the denaturation of the WT A*β*_42_ peptide and subsequent microsecond-length simulations using the Drude-2019 force field yielded an ensemble that was consistent with experimental properties and was subjected to further analysis. Given the consistency of the WT ensemble with experimental data, we applied the same protocol to the mutant peptides (see Methods).

### Clustering analysis

Dominant structures from each simulation ensemble were obtained by performing 2D-clustering in CHARMM. Thus, we were able to perform analysis on structurally similar subpopulations to relate their properties to the secondary and tertiary structures they manifested. Table 3 summarizes the outcomes of the clustering. The WT had the fewest clusters, while the simulations of mutant forms of A*β*_42_ yielded more clusters, reflecting more heterogeneous conformational ensembles. In contrast to the WT, E22K, and E22Q mutants, the E22G and D23N mutants both had numerous clusters with some only containing 1 snapshot (12 and 69, respectively).

**Table 3:**
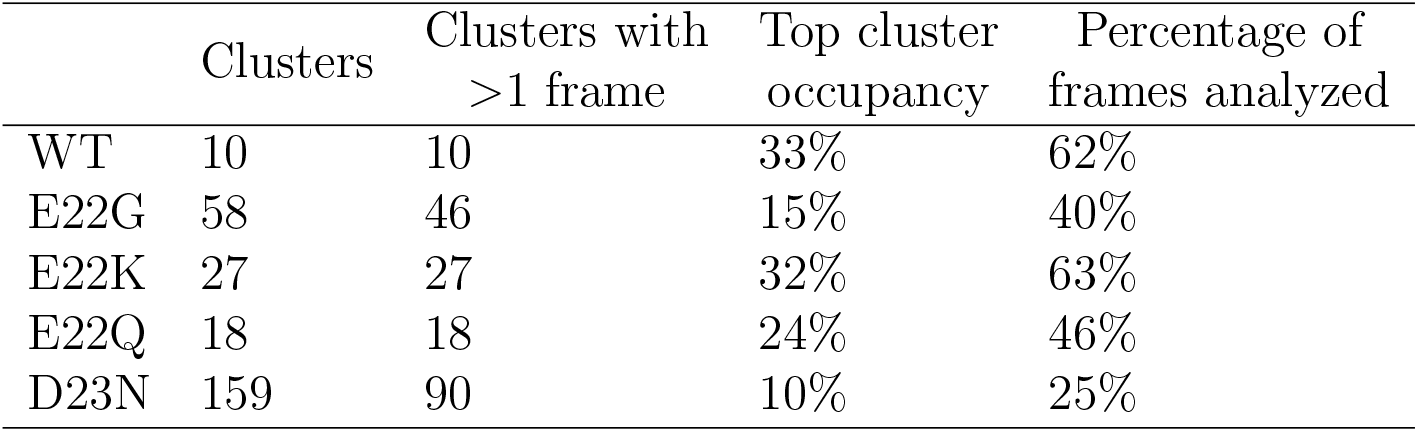
Statistics from 2D-clustering analysis

Given the considerable conformational heterogeneity of these peptides, we focused our analysis on the dominant conformations, defined as the top three clusters of each peptide ensemble, comprising 25-62% of the total snapshots (Table 3). The central (most representative) structure from the most-occupied cluster of each species is shown in Figure 2. The central structure for the second- and third-most-occupied clusters are shown in the Supporting Information, Figures S1 and S2. In the most-occupied cluster, we observed that the WT and E22K adopted a similar conformation in which the N- and C-termini interacted closely and formed a compact structure (Figure 2). Unlike the WT, the E22K mutant featured a short, antiparallel *β*-sheet involving N- and C-terminal residues. The E22Q and D23N mutants both lacked any defined structure and adopted extended, unfolded structures. The most-occupied cluster of the E22G ensemble was defined by a short *α*-helical turn but was otherwise mostly disordered.

**Figure 2:**
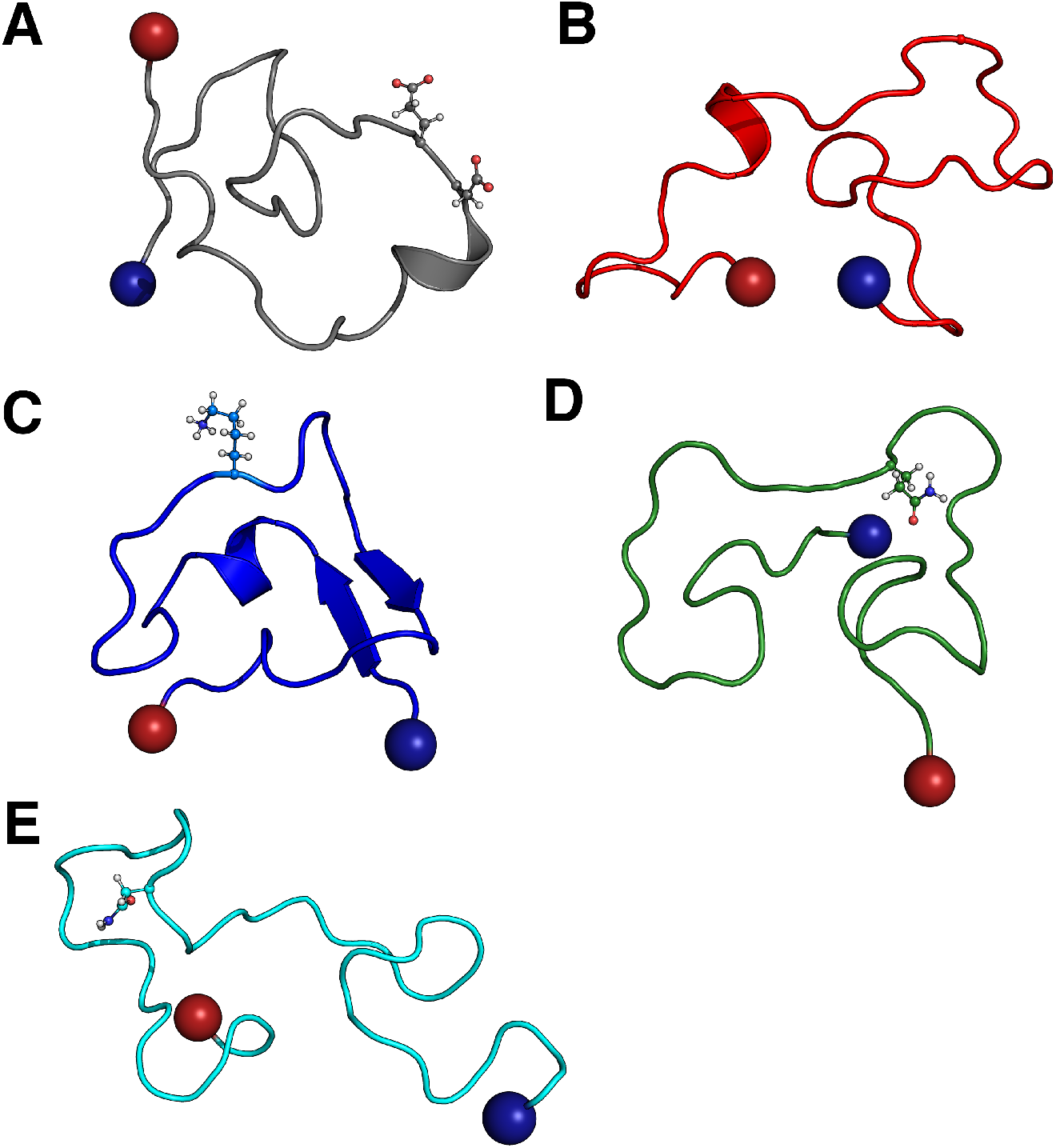
The central structure in each of the most-occupied clusters for (A) WT, (B) E22G, (C) E22K, (D) E22Q, and (E) D23N A*β*_42_ peptides. For perspective, residues 22 and 23 are depicted in ball and stick (both residues in the case of the WT and the relevant mutated residue in the other peptides). The N- and C-termini are indicated with blue and red spheres, respectively.

### Influence of mutations on secondary structure and hydrogen bonding network

To further characterize the structural characteristics of each peptide, we analyzed the secondary structure composition of each peptide in terms of *α*-helical and *β*-sheet occurrence at each residue in the sequence. In the most-occupied cluster of the WT, *α*-helicity was principally observed at residues Val24-Lys28, with less prominent *β*-sheet formation from Val12-Gln15 and Leu34-Gly37 (Figure 3). Mutations at position 22 led to shifts in *α*-helix location and/or an increase in *β*-sheet prevalence relative to the WT, suggesting that these mutations are somewhat ordering in terms of secondary structure. In the case of the E22G mutant, an *α*-helix persisted from residues Lys28-Ile32, shifted toward the C-terminus of the peptide than in the WT (Figure 3). The E22K mutant *α*-helix persisted toward the N-terminus and spanned residues Asp7-Val12 (Figure 3). Similarly, the position and likelihood of a *β*-sheet was perturbed by the E22K and E22Q mutations.

**Figure 3:**
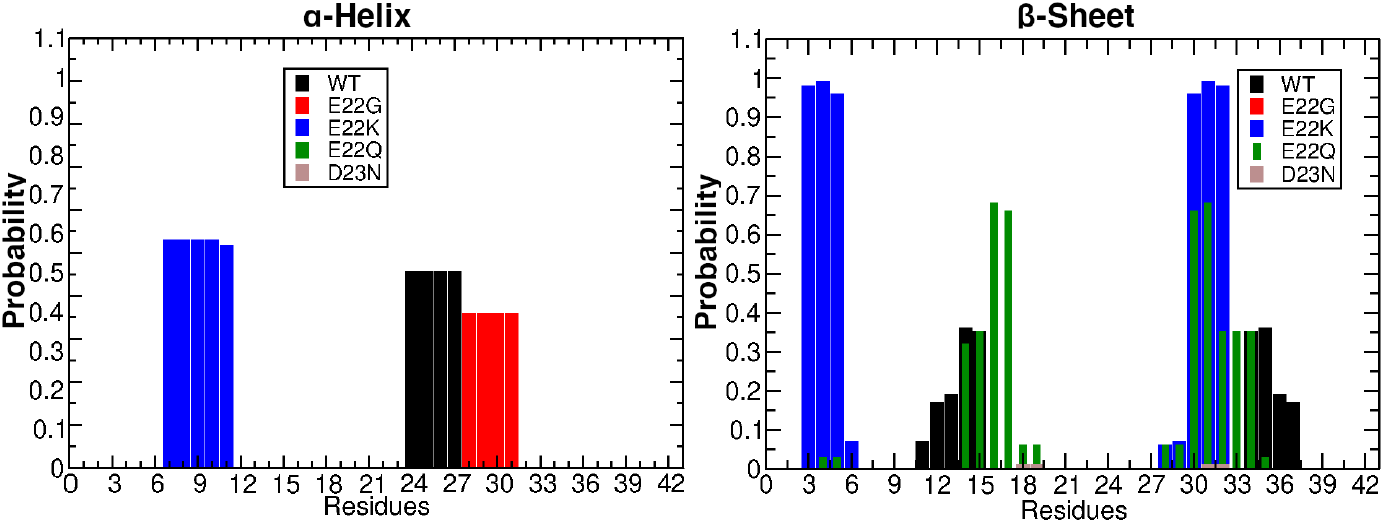
Per-residue probabilities of *α*-helix and *β*-sheet occurrence in the most-occupied cluster. Secondary structure types were assigned using DSSP^56^

In the WT, short *β*-sheets formed at residues Val12-Gln15 and Leu34-Gly37, though their occurrence was typically in less than 50% of the frames belonging to the most-occupied cluster. The E22K mutation led to a more pronounced *β*-sheet that included residues Glu3-Arg5 and Ala30-Ile32 (Figure 3), occurring in nearly all frames of the most-occupied cluster. These *β*-sheets were also present in the second-most-occupied cluster not the third-most-occupied cluster (Supporting Information, Figures S3 and SS4), suggesting some conformational heterogeneity but that this *β*-sheet is characteristic of the ensemble. Similarly, the E22Q mutant adopted a short *β*-sheet involving residues His14-Phe19 and Ala30-Leu34 (Figure 3). The formation of the *β*-sheet in the E22Q mutant persisted in the second-most-occupied cluster (Supporting Information, Figures S3 and SS4).

The D23N mutant was largely disordered, lacking *α*-helix or *β*-sheet structures (Figure 3), similar to the conformations adopted by the E22Q mutant. The nature of the E22Q and D23N mutants is such that the sidechain is altered from a carboxylate to an amide. We suspected that the lack of *α*-helical structure observed in the E22Q and D23N mutants arose from the formation of transient hydrogen bonds between the sidechain of Gln22 or Asn23 and the backbone amide groups. While transient, these hydrogen bonds existed closer to the N-terminus in the case of E22Q and toward the C-terminal residues in D23N (Supporting Information, Figure S5), and may have acted to disrupt the Val24-Lys28 *α*-helix that formed in the WT. These changes suggest that small perturbations in the charge and electronic structure of a single sidechain can alter the secondary structure within A*β*_42_.

The configurations adopted by the WT A*β*_42_ peptide in our simulations broadly agree with experimental descriptions of its structure, which have principally been obtained using NMR spectroscopy. An early study on the A*β*_12-28_ fragment characterized its structure as a “hydrophobically collapsed coil,” with the condensation of the CHC responsible for exposing other aggregation-prone residues.^71^ A subsequent investigation on the A*β*_10-35_ fragment and full-length A*β*_40_ found some secondary structure adoption, in the form of sharp turns indicated by NOE distances.^72^ The same study further concluded that Tyr10 and Val12 collapse onto the CHC residues to form a collapsed, hydrophobic structure. We observed these types of interactions in the WT peptide ensemble. The extent of residual secondary structure has recently been called into question by more recent NMR studies on A*β*_40_ and A*β*_42_, which have concluded that these peptides are essentially entirely unstructured, except for weak turn propensity around Asn27 and Lys28,^73^ and perhaps up to 10% *β*-strand in the C-terminal region.^74^

To further understand what role perturbations in electronic structure may have in driving changes in secondary structure, we calculated the peptide-bond dipole moments along the entire sequence (Supporting Information, Table S1). Alignment of peptide-bond dipole moments leads to cooperative induction that ultimately gives rise to the helix macrodipole, therefore changes in dipole moments may reflect changes in secondary structure and folding state.^37,65,75,76^ Changes in this property may also reflect solvent exposure of the polypeptide backbone. The sensitivity of peptide-bond dipole moments to structure and microenvironment allows us to investigate changes that may lead to mutation-specific alterations in the monomer ensemble that modulate A*β*_42_ aggregation pathways.

Per-residue peptide-bond dipole moments for all mutants were compared against those of the WT A*β*_42_ to determine regions of the polypeptide sequence that were most impacted by the mutations. Each of the mutants led to increased peptide-bond dipole moments in the CHC and C-terminal region (Figure 4). This increase in peptide-bond dipole moment may indicate a change in secondary structure (reflecting an altered hydrogen-bonding pattern) or a change in solvent accessibility. As will be discussed below, the increase in peptide-bond dipole moments arises from an increase in exposure to the aqueous solvent, which may have important consequences for these primarily hydrophobic sequences.

**Figure 4:**
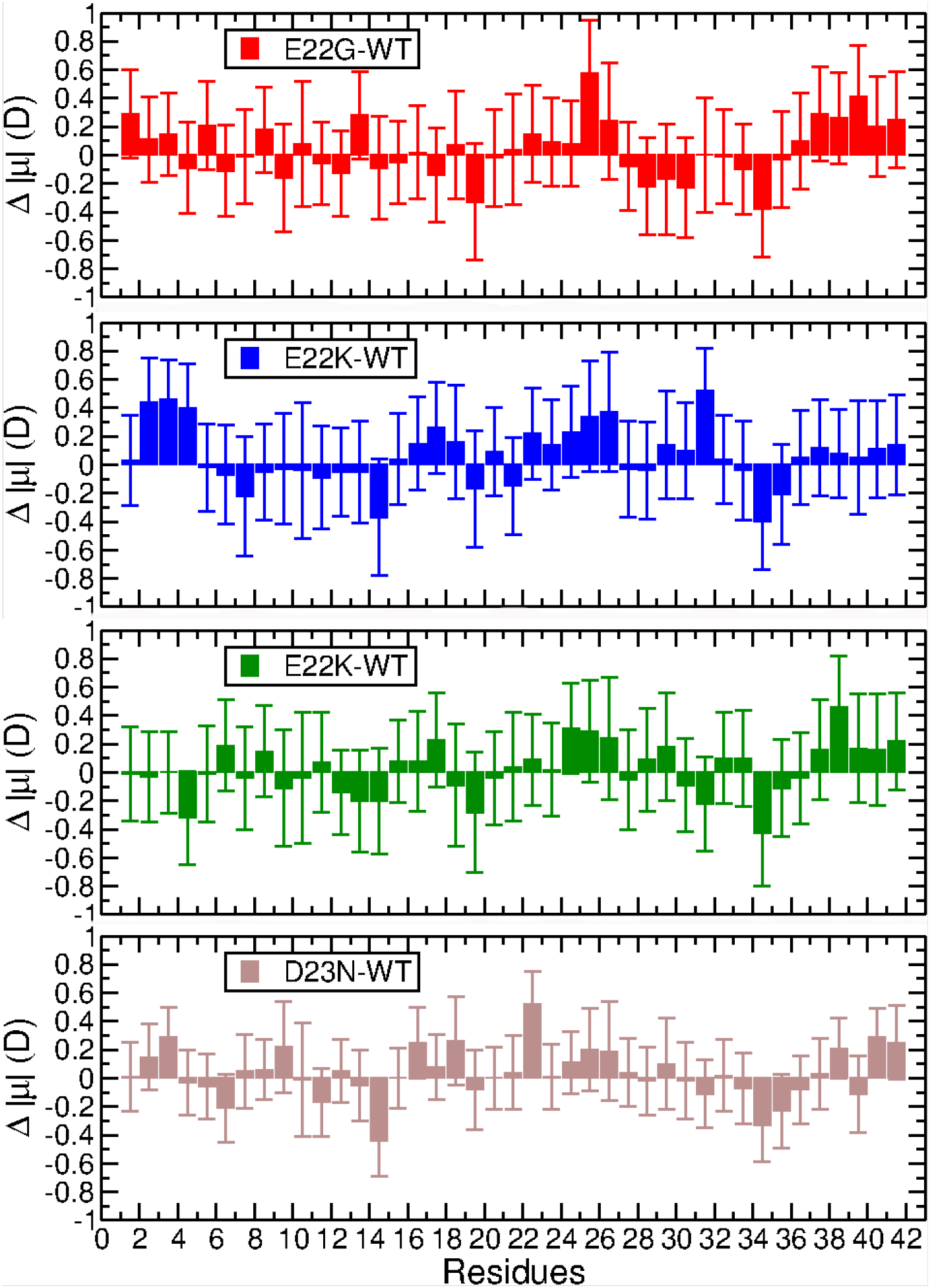
Difference in peptide-bond dipole moments relative to the WT. Error bars represent propagation of the standard deviation of each quantity.

We also observed that the peptide-bond dipole moments of residues surrounding Met35 were depolarized in each mutant relative to the WT. Ile32 in the E22K and E22Q mutants adopted a *β*-sheet structure, whereas this residue was disordered in the WT. Peptide-bond dipole moments in *β*-sheets are intrinsically lower than in random coils,^65^ which explains this difference. However, the E22G and D23N peptides lacked defined secondary structure around Met35, suggesting that the observed decrease in peptide-bond dipole was not driven by changes in secondary structure. In addition to the changes noted in the CHC and the hydrophobic C-terminal region, we observed an increase in Val24-Asn27 peptide-bond dipole moments in the mutants (Figure 4 and Supporting Information, Table S1). This increase in dipole moment was due to the lack of *α*-helical secondary structure in this region in the mutants. A previous study showed that peptide-bond dipole moments are larger in random coils than those of *α*-helices.^65^ Thus, the preference for disordered states among residues Val24-Asn27 in the mutants is reflected in the dipole moments. These trends were also present in other clusters (Supporting Information, Figure S6), suggesting a prominent role for polarization of individual peptide bonds in defining the properties of the overall conformational ensembles.

The absolute value of the peptide-bond dipole moments is influenced by the intrinsic geometry of the polypeptide backbone, so to determine the contributions of inductive effects due to solvation and sidechain properties, we calculated the peptide-bond helical dipole enhancement of the mutants and WT using the method described by Huang and MacKerell.^65^ Computing this quantity for helical structures is critical for understanding the helix-coil transition that underlies many of the conformational changes that A*β*_42_ undergoes in its initial unfolding events. Because there was little overall *α*-helicity in the top three clusters, we calculated the dipole enhancement using the whole trajectory (3 *μ*s) to sample this quantity more comprehensively. Enhancements were subsequently compared those of WT A*β*_42_.

In general, the peptide-bond helical dipole enhancements decreased for residues immediately surrounding each mutated residue (Figure 5). This decrease in enhancement suggests that the hydrogen bonding network surrounding the mutation is weakened in the mutants, explaining why we observed little *α*-helicity in the most-occupied clusters. In addition, the E22K, E22Q, and D23N mutants had enhanced peptide-bond dipole moments from residues Gly33-Gly37, a hydrophobic region near the C-terminus (Figure 5). This dipole enhancement indicates that the backbone amide groups of these hydrophobic residues engaged in more favorable hydrogen bonding in the mutant structures than in the WT. These results suggest that the other attributes of the mutants, such as the increased dipole moments that favor solvation described above, destabilize the hydrogen bonding network that would otherwise be required for maintaining helical structure.

**Figure 5:**
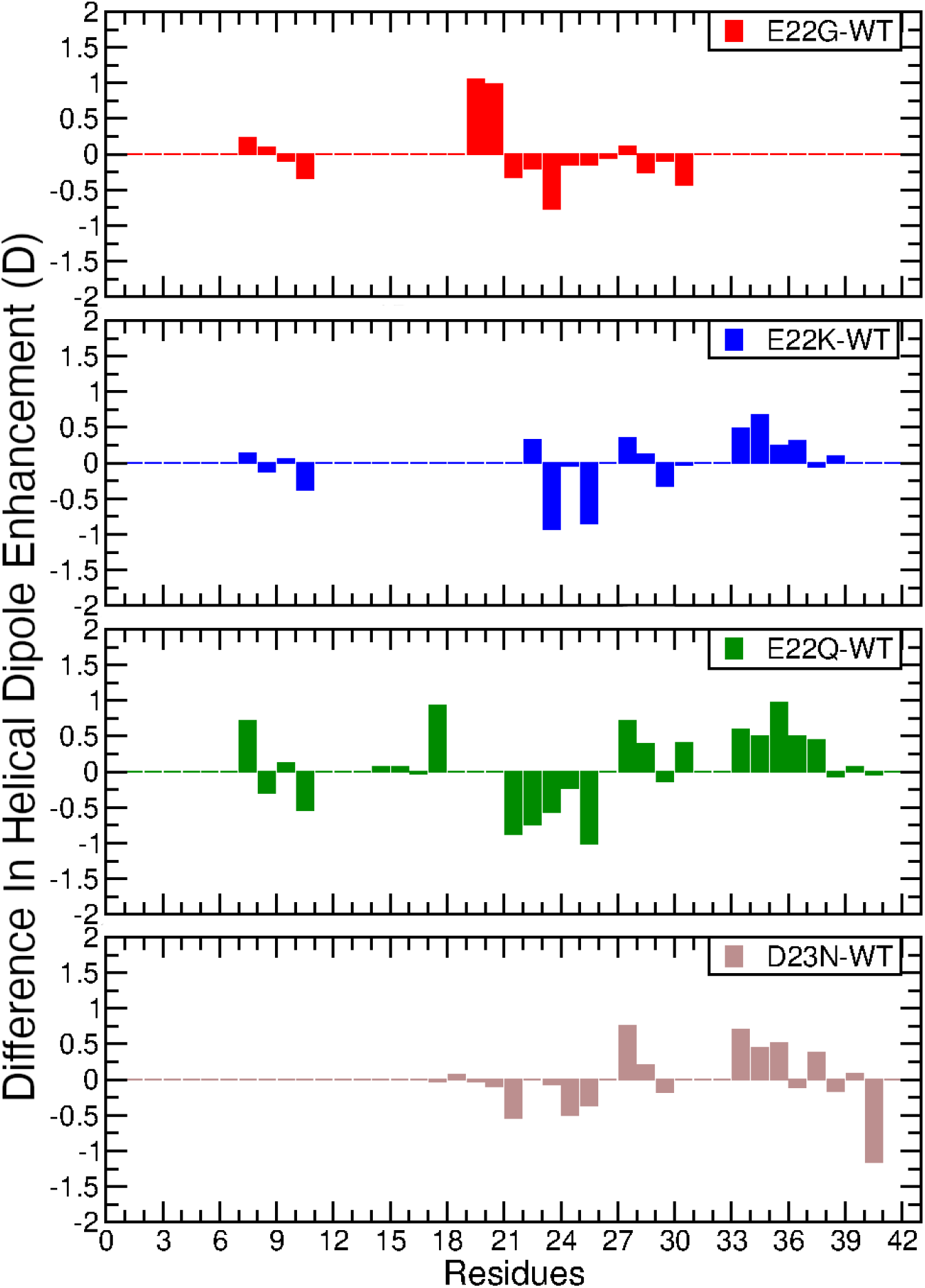
Difference in helical dipole enhancement of the full-length simulations compared to the WT. Zero values indicate an absence of *α*-helical structure at that residue for either the WT or mutants.

There are some notable differences among the helical dipole enhancements of the mutants. In the E22G mutant, the helical peptide-bond dipole enhancements between Phe19-Phe20 and Phe20-Ala21 were greater than in the WT, suggesting stronger induction when in an *α*-helix. However, *α*-helical structures were absent in the most-occupied clusters, suggesting that other aspects of the E22G mutant destabilize an otherwise stable *α*-helix in a similar way that the E22K, E22Q, and D23N mutant destabilized the hydrogen bonding network of the C-terminal residues. Together, these results suggest that although each mutant may transiently form *α*-helical structures, in some cases exhibiting strong peptide-bond dipole enhancements that would stabilize *α*-helices, other aspects of the mutants destabilize these hydrogen bonding networks. Below, we describe how tertiary interactions and perturbations in sidechain dipole moments may contribute to this phenomenon.

### Impact of tertiary hydrophobic interactions on A*β*_42_

Having assessed the relationship between secondary structure content and induced dipoles in the polypeptide backbone, we sought to characterize the tertiary structure of each A*β*_42_ peptide. Ultimately, the contributions of secondary structure (disorder) and tertiary structure (compactness and accessibility of aggregation-prone sequences) will govern the earliest events of oligomer nucleation.

Using the most-occupied cluster, we calculated the pairwise distances between all C*α* atoms as a way to describe residue-residue contacts (Figure 6). In addition to the C*α* distances, we calculated the distance matrix for all heavy atoms to understand the nature of the contacts observed in the C*α* distance matrix (Supporting Information, Figure S6). The WT heavy-atom distance matrix was subtracted from those of the mutants to generate difference distance matrices (Supporting Information, Figure S9), which can be used to visualize shifts in interatomic contacts. We subsequently computed the solvent-accessible surface area (SASA) of each residue (Supporting Information, Table S2 and Figure 7) to determine how shifts in interatomic contacts impacted exposure of aggregation-prone regions of the peptides. Thus, we focus our discussion here on shifts in tertiary structure of each mutant relative to the WT.

**Figure 6:**
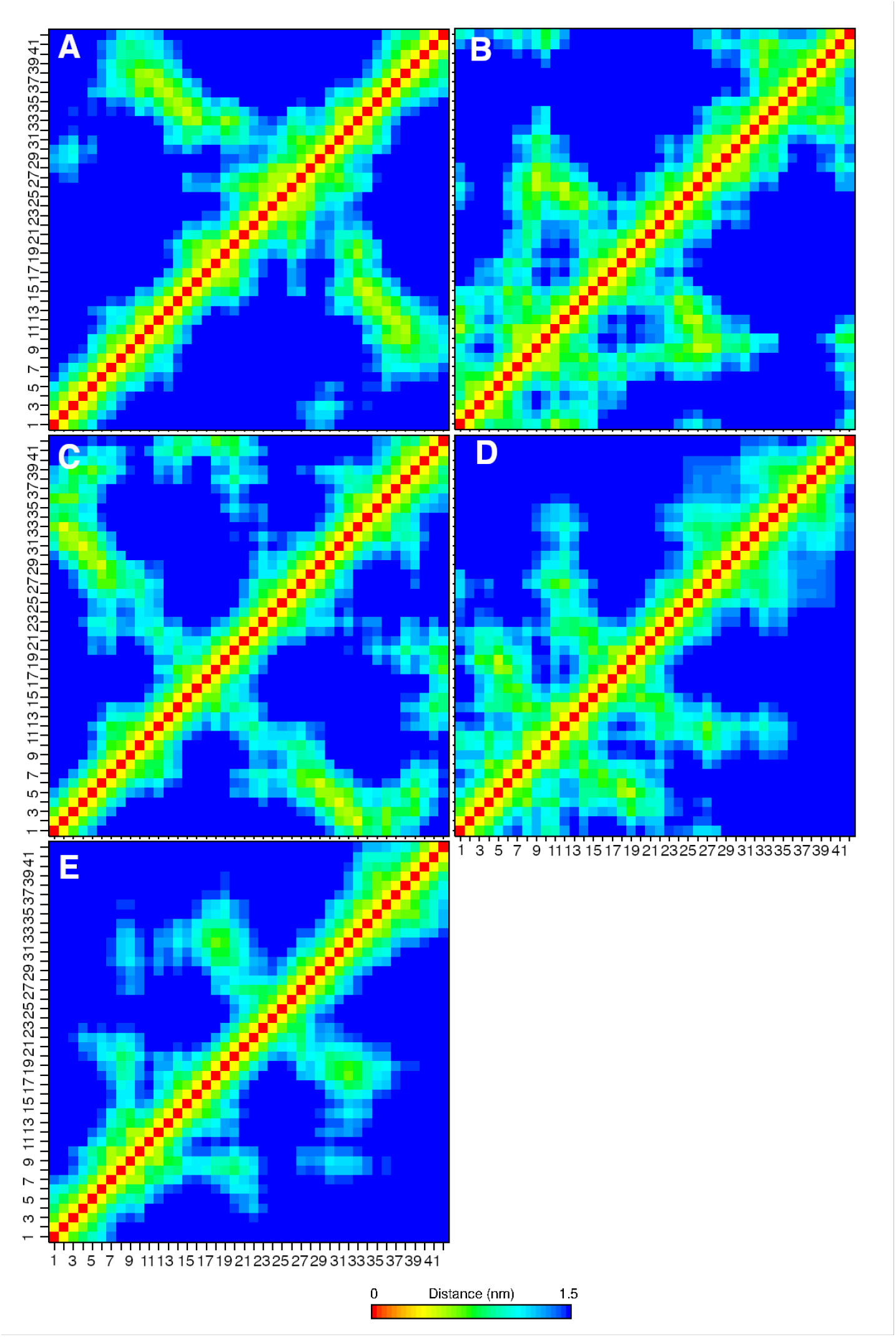
C*α*-C*α* distance matrix of the most-occupied cluster used to calculate tertiary structure interactions for the (A) WT, (B) E22G, (C) E22K, (D) E22Q, and (E) D23N peptides.

**Figure 7:**
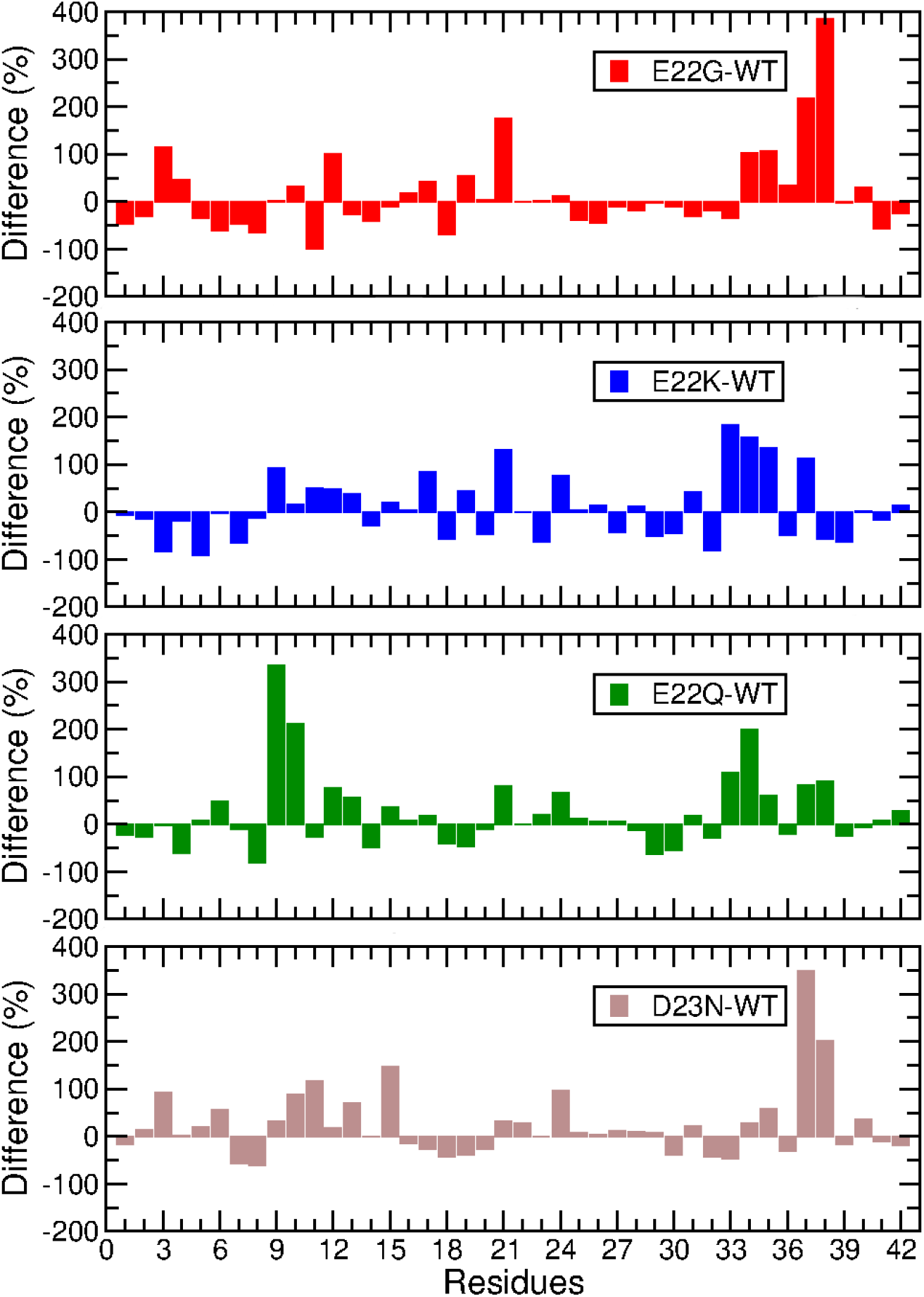
Relative change in solvent-accessible surface area (SASA), expressed as a percentage relative to the WT values. Data were collected for structures belonging to the most-occupied cluster from each simulation set.

The N-terminal residues of the E22G mutant interacted closely with the CHC and the residues surrounding Gly22 (Supporting Information, Figures S6B and SS9A). In addition, residues Gly25-Leu34 had close contacts with the C-terminus. This shift in contacts led to dramatic increases in SASA for residues Glu3, Val12, Ala21, and Leu34-Gly38 (Figure 7). In contrast, Glu11 had a marked decrease in SASA. The simultaneous exposure and shielding of Glu3 and Glu11, respectively, likely have no net impact on the solubility of the E22G peptide, whereas the other residues with increased SASA are hydrophobic. The increased SASA of Ala21 may a result of the change of Glu22 to Gly22, which neutralizes and removes the sidechain, allowing water to interact more closely with Ala21. The increase in solvent exposure at the aggregation-prone C-terminus may contribute to the enhanced aggregation rate of the E22G peptide.

The E22K mutant adopted a similar tertiary structure to the WT in which the N- and C-termini folded onto themselves (Figure 2A,C), leading to reduced SASA for N-terminal residues and the region encompassing Lys28-Ile32 (Figure 7). However, we observed differences between the two when we compared the heavy-atom distance matrixes (Supporting Information, Figure S9B). Like the E22G mutant, the N-terminal residues of the E22K mutant also interacted closely with the CHC, but these interactions mainly involved backbone atoms in the CHC. Another factor in the decreased SASA around Lys28-Ile32 was the formation of a two-stranded *β*-sheet encompassing these residues, suggesting packing of the sidechains that may shield them from the aqueous solvent. As with the E22G mutant, several residues in the hydrophobic C-terminal region became more solvent-exposed (Gly33, Leu34, Met35, and Gly37) relative to the WT (Figure 7), suggesting the possibility for enhanced aggregation in this critical region. The Ala2-His6 sequence interacted closely with Gly29-Gly33, driven by the sidechain of Glu3 forming hydrogen bonds with the backbone of Gly37 and Gly38, interactions that may have promoted the solvent exposure of nearby sidechains.

The interatomic contacts of the E22Q mutant reflected a more compact N-terminal region that lacked some of the longer-range contacts observed in the WT (Figure 6 and Supporting Information, Figure S9C). One specific cluster of long-range interactions observed in the E22Q mutant was between Asp1-His6 with Ile31-Val36. In addition, Phe4 formed close contacts with Asp1 and Ser8 (Supporting Information, Figure S9C), which we will discuss in more detail later in the context of its shifted sidechain dipole moment. The E22Q mutant had a pattern of solvent accessibility that largely mirrored that of the E22K mutant, with the exception of the prominent increase at Gly9 and Tyr10 (Figure 7). This change in solvent accessibility may be caused by the shift in location of the *β*-sheets that formed in the WT and E22Q. In the WT, Tyr10 was shielded by Arg5 and His13, resulting in a less solvent-exposed Gly9 and Tyr10. As shown above, the *β*-sheets of the WT formed at residues Val12-Gln15 and Leu34-Gly37, whereas the E22Q *β*-sheets spanned His14-Phe19 and Ala30-Leu34. The shift in secondary structure in the E22Q mutant caused Gly9 and Tyr10 to be less shielded by the residues that comprised the WT *β*-sheets. Similar to the E22G and E22K mutants, the E22Q mutant had increased SASA among C-terminal residues, specifically Gly33, Leu34, and Met35.

The D23N mutant had the most elongated structure among the variants examined here, forming mostly short-range contacts. However, the D23N mutant had a few sidechain interactions between Phe19 and hydrophobic C-terminal residues Gly33-Ala42 (Supporting Information, Figure S9D). The adoption of primarily short-range contacts led to the D23N mutant manifesting similar solvent accessibility to the E22Q, marked by increases among N- and C-terminal residues (Figure 7). The shared patterns of relative increases and decreases in SASA by these chemically similar mutations (both E22Q and D23N replace an acid with an amide) suggest they may impart similar properties to the A*β*_42_ peptides.

Across all the mutants examined here, the solvent accessibility of Gly37 and/or Gly38 was increased (Supporting Information, Figure S7), a phenomenon that was also related to an increase in the peptide-bond dipole moments for these residues (Figure 4). Shifts in peptide-bond dipole moments may be related to changes in secondary structure (via induction through hydrogen bonds) or solvent exposure. The observed increases in Gly37 and Gly38 dipole moments were not related to any specific change in secondary structure, instead, they were associated with an increase in solvent accessibility. The increase in peptide-bond dipole moments indicates that these hydrophobic regions are more polarized in the mutants, allowing them to be more solvent-exposed. Previous work has shown that during amyloid formation, Met35 packs against Gly37, creating a molecular notch that promotes fibril formation.^77,78^ Our observed increase in solvent accessibility in the mutants, as well as increases in peptide-bond dipole moment, suggest that perturbations in the properties of these residues also affect the conformational sampling of this critical region in A*β*_42_ monomers.

### Sidechain dipole moments highlight changes in solvation and heterogenous microenvironments

Once we had a general understanding of the tertiary structure of the WT and mutants, we investigated how the dipole-dipole interactions changed as a function of structure, which we expected would reflect unique microenvironments. As such, we calculated the sidechain dipole moments and compared them to the WT (Figure 8), omitting direct comparisons between the mutated residues. That is, changing the chemical nature of a given sidechain imparts a completely different dipole moment, whereas our interests were in systematically evaluating how mutations perturbed the electronic structure of the remainder of the A*β*_42_ peptide. In this analysis, three regions of the A*β*_42_ peptides emerged as having distinct dipole responses: (1) Phe4 and surrounding residues, (2) the CHC, and (3) the hydrophobic C-terminal region.

**Figure 8:**
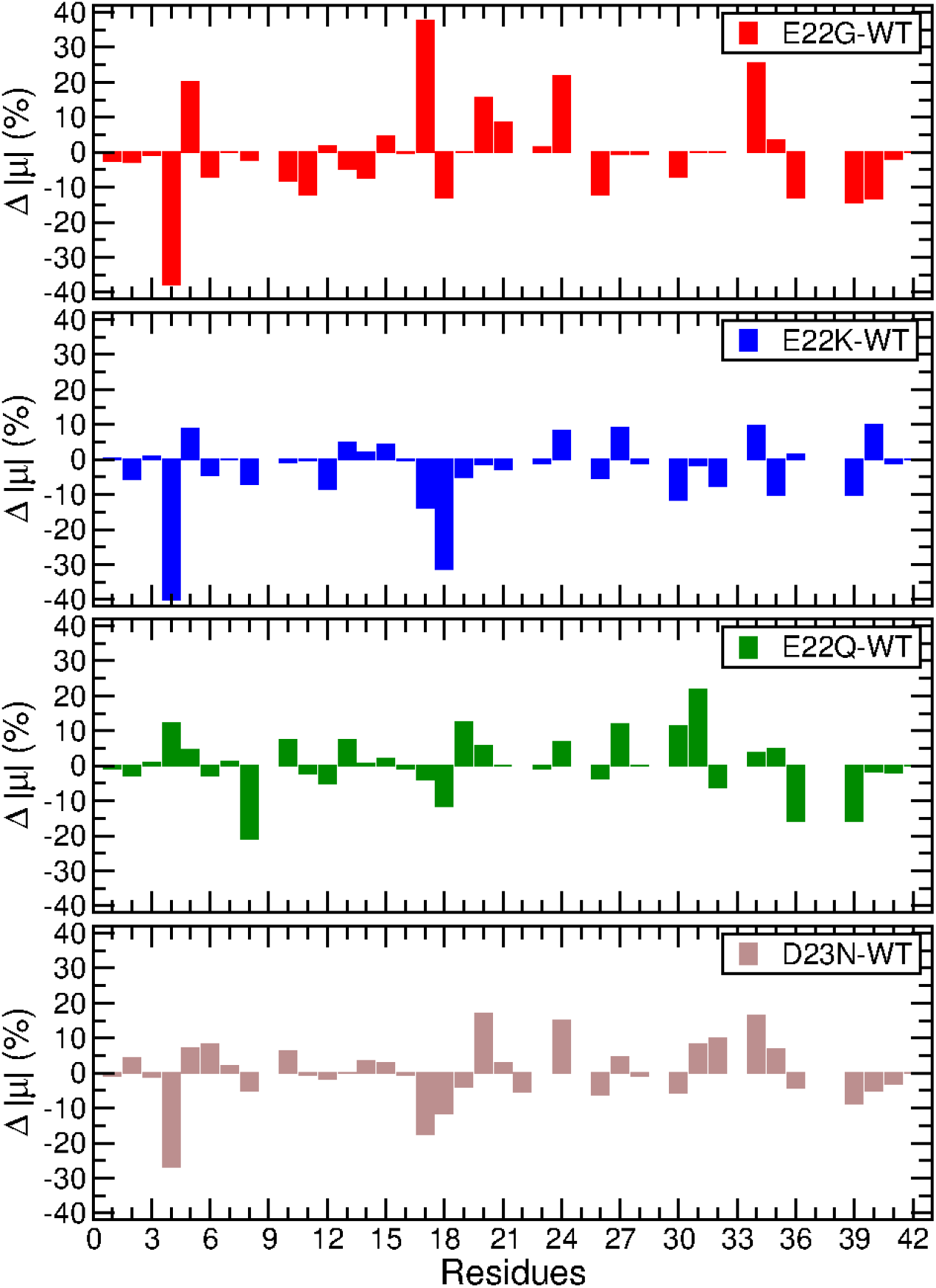
Percent change in sidechain dipole moment, expressed as a percentage relative to the values of the WT A*β*_42_ peptide. Mutated residues were omitted from this analysis, as were glycine residues, as they have no sidechain atoms.

Phe4 was strongly depolarized in the E22G (sidechain dipole moment of 0.8 ± 0.3 D), E22K (0.8 ± 0.3 D), and D23N (0.9 ± 0.4 D) mutants relative to the WT (1.3 ± 0.4 D) peptide (Supporting Information, Table S3). We hypothesize that the WT Phe4 was more polarized than the mutants because it was more solvent-exposed than the E22K and D23N mutants and engaged in interactions with the sidechain of Asp1, which the mutants did not. Phe4 in the E22Q mutant was slightly more polarized than the WT, with a sidechain dipole moment of 1.5 ± 0.4 D. Phe4 of E22Q was less solvent-accessible than in the WT, surrounded by Asp1, Tyr10, and the backbone of Ala2. The proximity of Asp1 and Tyr10 in this occluded microenvironment likely led to the increased sidechain dipole moment of Phe4 in this mutant. These changes in dipole moments highlight the sensitivity of the Phe4 to its microenvironment.

The CHC harbors the mutations that were made in each peptide, therefore we anticipated that this region would have perturbed electrostatic properties as a result. However, the CHC experienced very few perturbations in sidechain dipole moments (Figure 8 and Supporting Information, Table S3). One of the major changes was that of the Leu17 in the E22G mutant, which had an increased sidechain dipole compared to the WT (Figure 8). We found that Leu17 interacted closely with His6 and His14, forming an unusual microenvironment with both polar and nonpolar residues. His6 and His14 both depolarized relative to the WT values (Supporting Information, Table S3), highlighting the plasticity of dipole moments and their ability to shift in response to heterogeneous interactions. As a result, many other residues in the polar N-terminal region of the E22G mutant had depolarized sidechains, resulting in an N-terminus that is slightly more hydrophobic, which could also explain why the E22G mutant is more aggregation-prone, in addition to the increased hydrophobic C-terminal SASA described above.

Other important shifts in the sidechain dipole moment occurred within the hydrophobic C-terminal region. In the E22G, E22Q, and D23N mutants, Val39-Ala42 had decreased sidechain dipole moments relative to the WT (Figure 8). E22K was similar to the other mutants but Ile41 had an increased dipole moment, which may have been due to the proximity of the sidechains of Gln15 and Lys16 (Supporting Information, Figure S9 and Table S3). This observation is another example of a heterogeneous microenvironment that may be relevant in dictating the conformational ensemble of the E22K monomer. Across the mutants, the combination of polarization of the backbone amide groups that will favor solvation (Figure 4), depolarization of the hydrophobic sidechains (Figure 8), and increased solvent accessibility of residues such as Gly37 and Gly38, suggest that the mutants cause the hydrophobic C-terminal region to be more aggregation-prone.

### Thermodynamic properties of A*β*_15-27_

To gain a further understanding of how the mutations of A*β* alter the folding thermodynamics of the protein, we carried out US simulations to calculate the ΔG_fold_. While A*β* is not believed to adopt a stable *α*-helix in the region encompassing the CHC in aqueous solution, here we sought to understand the impact of mutations on the nature of the helix-coil equilibrium. That is, to what extent does mutation (de)stabilize an *α*-helical structure and affect the likelihood of disorder in this key region? We previously demonstrated that FAD mutations impact sidechain dipole properties and unfolding pathways of this fragment in unbiased simulations,^37^ and here we sought to augment our understanding of the properties of this key region. We performed this analysis over a range of temperatures to resolve the contributions of entropy and enthalpy to this quantity. We used a peptide fragment of A*β* encompassing residues 15-27 to systematically analyze how the mutations altered ΔG_fold_ and determine how dipole-dipole interactions in the CHC were perturbed in response to the changes in charge. As described in the Methods, we performed 100-150 ns of simulation time in each of 32 windows for each variant, and only analyzed the final 50 ns after determining convergence of ΔG_fold_ from 5-ns windows of time. The results of these calculations are provided in the Supporting Information, Figure S10 and Table S4.

The resulting free energy profiles had similar features, with minor differences around the two minima in the range of 18 – 21 Å (Figure 9 and Supporting Information, Figure S11). The WT and E22G had global minimum at ∼20.5 Å, whereas the E22K, E22Q, and D23N mutants had global minima around ∼18.5 Å (Figure 9), indicating that the WT prefers to be slightly elongated compared to these mutants. The free energy profile showed that the E22G variant had the smallest difference between the two minima, which may be due to the flexibility of Gly22 at the center of the peptide facilitating the interconversion between these two states.

**Figure 9:**
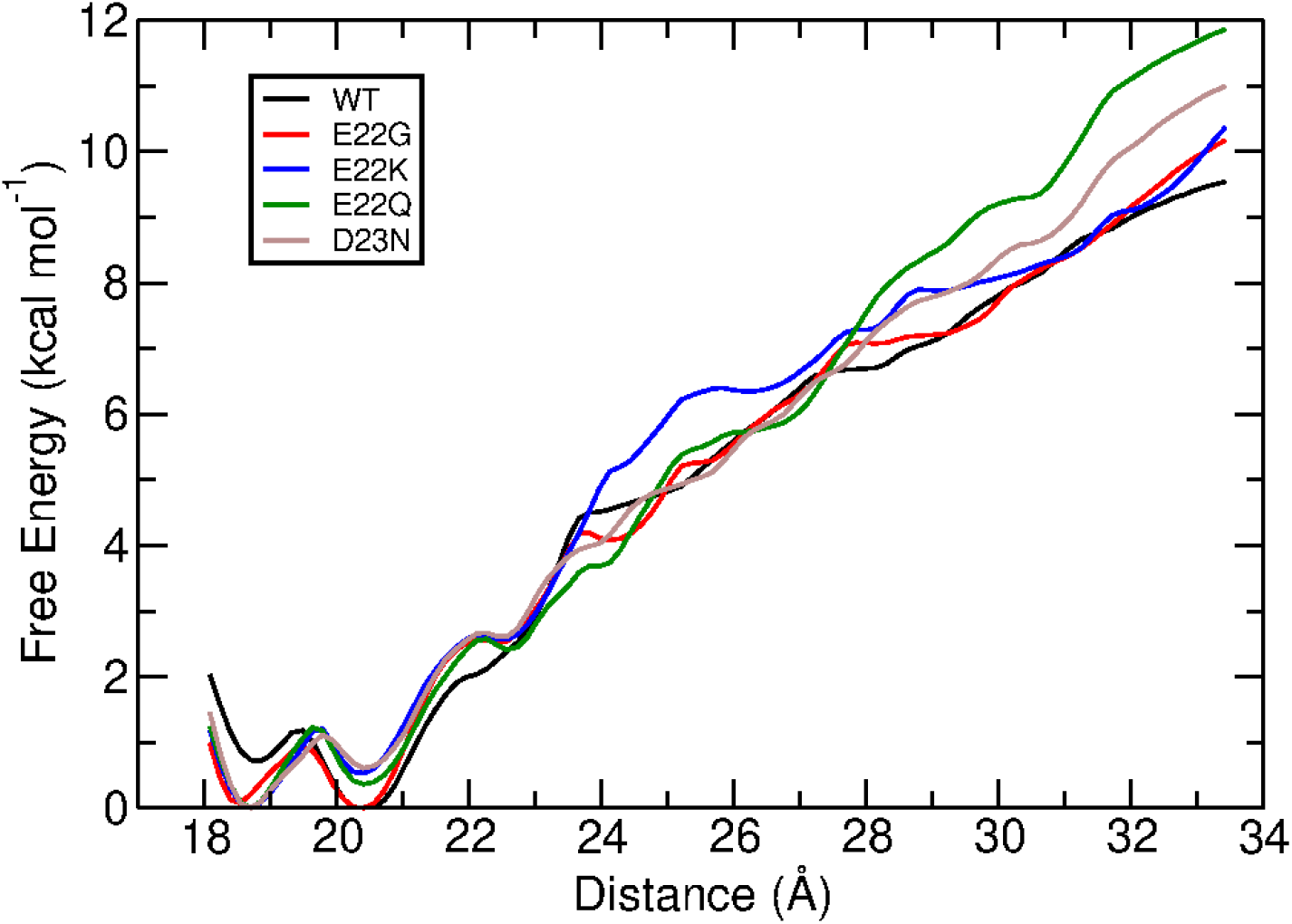
Free energy profiles at 298 K for the different A*β*_15-27_ variants, calculated by WHAM.

From the WHAM-derived free energy profiles, we calculated ΔG_fold_ at 298 K by summing over the ensemble of folded and unfolded states (see definition in Methods). We found that the WT had a folding free energy of -4.6 ± 0.7 kcal mol^-1^, whereas the E22 mutants all had more favorable ΔG_fold_ than the WT (Table 4). The E22G mutant had the least favorable ΔG_fold_ among the E22 mutants, which may be due to the E22G mutant having a free energy surface that most closely resembled that of the WT. The D23N mutant, however, had the least favorable ΔG_fold_ among all the variants investigated here, -3.6 ± 0.8 kcal mol^-1^ (Table 4).

**Table 4:** ΔG_fold_ (kcal mol^-1^) of the A*β*_15-27_ variants from US simulations at 298 K.

| | $\Delta G_{\text{fold}}$ |
| --- | --- |
| WT | $-4.6 \pm 0.7$ |
| E22G | $-5.4 \pm 0.3$ |
| E22K | $-6.1 \pm 0.6$ |
| E22Q | $-5.6 \pm 0.8$ |
| D23N | $-3.6 \pm 0.8$ |

Given that helix-coil equilibrium is driven to some extent by cooperativity of induced dipoles along the polypeptide backbone,^75^ we calculated helical dipole moment enhancements for each peptide bond in the peptide. The alignment of these hydrogen bonds enhances the peptide-bond dipole moment during helix nucleation.^76^ The enhancement thus allows us to track the formation and breakage of hydrogen bonds and evolution of secondary structure throughout the US simulations in terms of induced dipoles. Specifically, we computed the peptide-bond dipole moment enhancement for residues that satisfied the *α*-helical criteria (as defined by Huang and MacKerell)^65^ to see the dipole response in each variant as its secondary structure was lost as a function of extension along the reaction coordinate.

Unfolding of the WT, E22G, E22K, and E22Q all proceeded similarly, beginning at the N-terminus, as reflected by the degradation in dipole enhancement corresponding to loss of cooperativity in the backbone hydrogen-bonding network (Figure 10). The unfolding pathway taken by the WT and E22 mutants is likely the reason for the similar ΔG_fold_ values in these variants (Table 4). The E22K mutant had the most favorable ΔG_fold_, which could not be explained solely by these dipole enhancements. As was observed previously in unbiased simulations of this mutant, the formation of a salt bridge between Lys22 and Asp23 perturbed the properties of this variant.^37^ From 18-21 Å along the reaction coordinate, the Lys22-Asp23 salt bridge formed between 7-25% of the simulation time (Supporting Information, Figure S12). As the peptide unfolded further (between 26-33.5 Å along the reaction coordinate), this salt bridge formed more frequently, between 15-37% of the time (Supporting Information, Figure S12). This shift in salt-bridge stability suggests that favorable solvation of Lys22 and Asp23, leading to salt bridge disruption, promotes the adoption of helical states.

**Figure 10:**
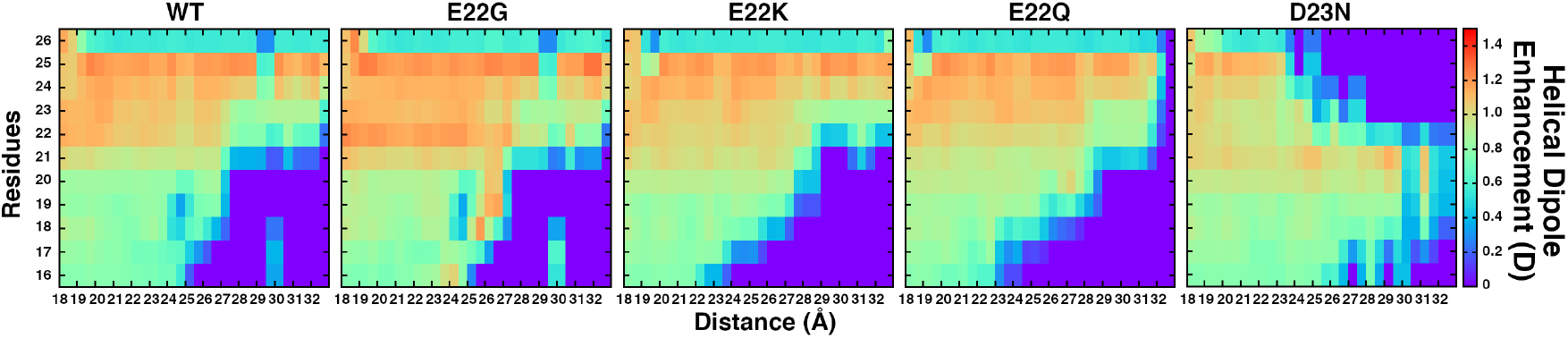
Helical dipole enhancement values (D) over the last 50 ns of the US simulations performed at 298 K.

**Figure 11:**
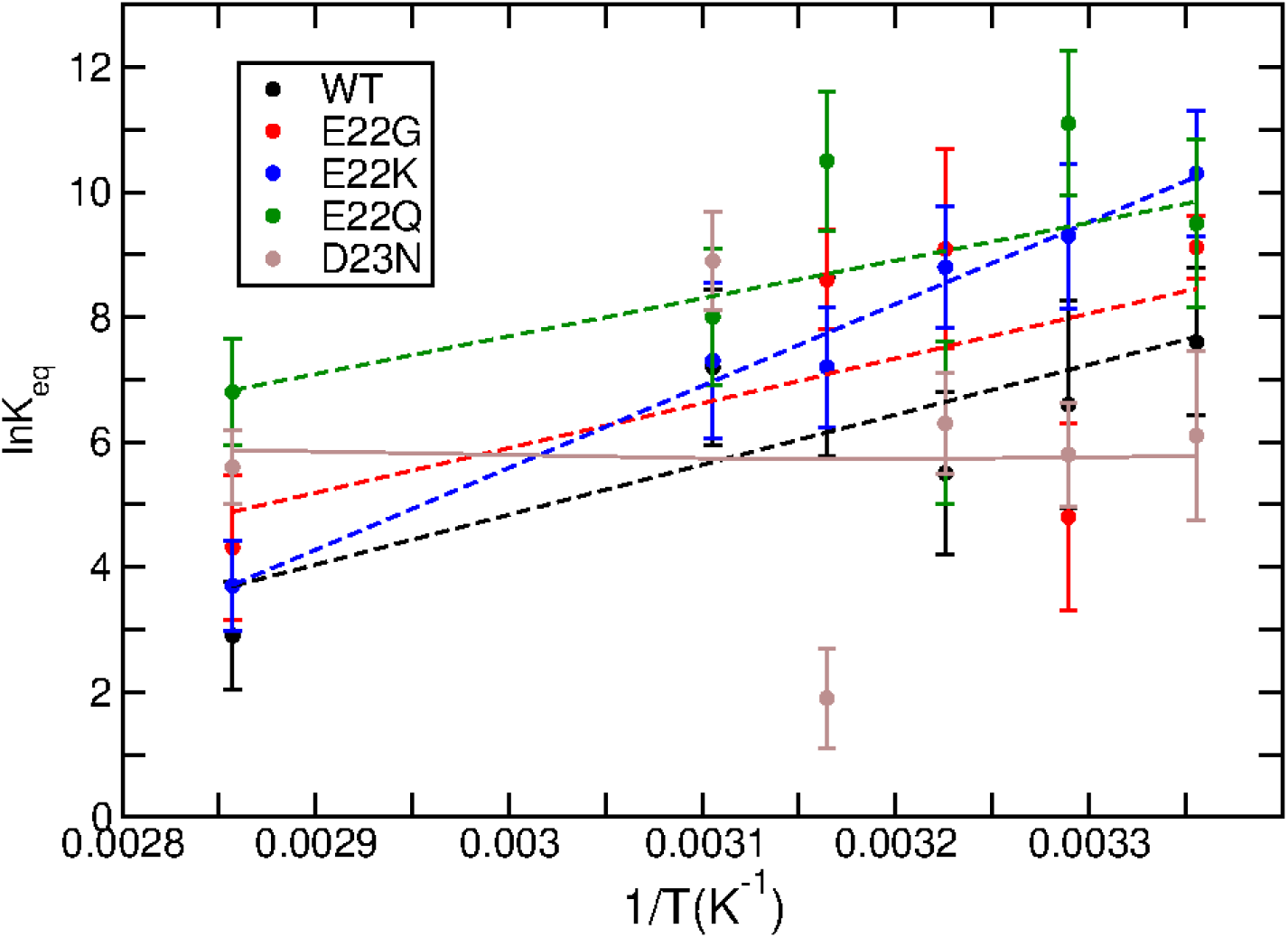
van’t Hoff analysis from US simulations performed over a range of temperatures. Dashed lines indicate systems for which the linear van’t Hoff equation was applied, and solid line indicates the application of the quadratic form of the van’t Hoff equation (D23N only).

The D23N variant unfolded differently from the WT and E22 mutants, unfolding from the C-terminus, followed by N-terminal residues subsequently disordering around 26 Å along the reaction coordinate (Figure 10 and Supporting Information, Figure S13). The different unfolding pathway, which led to simultaneous degradation of peptide-bond dipole enhancement from both ends of the A*β*_15-27_ peptide, explains why the D23N has the least favorable folding free energy. The mutation weakens the hydrogen-bonding network at both termini of the peptide.

To decompose the ΔG_fold_ values into ΔH and TΔS contributions, we repeated the US simulations at 304, 310, 316, 322, and 350 K, allowing us to perform a van’t Hoff analysis following conversion of ΔG_fold_ to lnK_eq_. This analysis revealed that the WT A*β*_15-27_ peptide and the E22G, E22K, and E22Q mutants all follow the linear van’t Hoff relation (Eq. 2). From this linear equation, we determined that the WT, E22G, E22K, and E22Q peptides had enthalpic and entropic contributions that both contributed favorably to ΔG_fold_, with ΔH dominating (Table 5). Furthermore, we see that the E22K mutant has the most favorable ΔH, which could be caused by the solvation of Lys22 and Asp23 in the *α*-helical state.

**Table 5:** ΔH and TΔS (both in kcal mol^-1^) contributions to ΔG_fold_ for all A*β*_15-27_ variants.

| | $\Delta H$ | $T\Delta S$ |
| --- | --- | --- |
| WT | -15.9 | 11.3 |
| E22G | -14.2 | 9.2 |
| E22K | -26.0 | 20.0 |
| E22Q | -12.0 | 6.2 |
| D23N | -1.0 | 2.4 |

The D23N peptide proceeded along a different unfolding pathway (Figure 10), suggesting a more complicated description of its equilibrium, one in which a simple two-state model may fail. Thus, the linear van’t Hoff equations could not be used to calculate ΔH and TΔS. Instead, we applied the quadratic form of the van’t Hoff relationship, finding that the D23N mutant is entropically driven, as it is the only peptide for which the magnitude of TΔS larger than that of ΔH. This difference may be caused from the D23N peptide unfolding from both the N- and C-termini. This result also rationalizes why the full-length protein unfolds and remained the most disordered of all the variants investigated here.

## Conclusions

Here, we have performed the first systematic study of the full-length monomeric WT A*β*_42_ and pathogenic, familial AD mutations using a polarizable force field. Additionally, we investigated the monomer thermodynamic properties of the A*β*_15-27_ fragment and its mutants. We sought to understand the initial unfolding events and conformational changes of the fulllength WT monomer and mutants that are known to be more aggregation-prone. We found that the mutations at position 22 resulted in shifts in *α*-helix location and/or an increase in *β*-sheet prevalence relative to the WT. The D23N mutated resulted in a complete loss of *α*-helix and *β*-sheet structure.

Perturbations in peptide-bond dipole moments reflected shifts in secondary structure or increased favorability of solvation. Mutated forms of A*β*_42_ generally had elevated peptidebond dipole moments in the hydrophobic C-terminal region relative to the WT peptide, making this region more amenable to solvation. To account for intrinsically different peptide-bond dipole moment values across secondary structure types, we computed helical dipole enhancements and found that mutations at positions 22 and 23 weakened helical structures in their vicinity. Together, these results suggest that although each mutant may transiently form *α*-helical structures, the dominant structures result from destabilized the hydrogen bonding networks, leading to shifts in secondary structure within the mutants. As a result, FAD mutants had increased SASA at the hydrophobic C-terminal region, which may have a role in fibril formation.

Sidechain dipole moments may also play a role in dictating the conformational transitions of A*β*_42_ and accommodating heterogeneous microenvironments. Phe4 was notable in this regard, as it exhibited shifts in its sidechain dipole moment in response to interactions with polar groups, but Phe19 and Phe20, which reside near the sites of mutation, were not impacted, and overall lack of sidechain dipole moment response in the CHC was surprising, though Leu17 exhibited some shifts in response to atypical interactions with polar residues. However, the lack of sidechain dipole moment changes suggests that although the mutations may change the chemical nature of a given sidechain, the mutations alter the CHC through changes in the peptide-bond dipole moments and other long-range contacts that lead to different tertiary structures and therefore various contacts and microenvironments.

Using US to unfold A*β*_15-27_, which contains the mutation sites and the CHC, we found that WT and E22 mutant folding is enthalpically driven, whereas D23N folding is entropically driven. In addition, the D23N mutant underwent a different unfolding pattern than the other mutants. Taken together, the full-length and fragment simulations demonstrate that polarization of the backbone and depolarization of sidechains play a pivotal role in dictating the conformational ensemble of A*β*_42_ and perhaps the initial unfolding events along the amyloidogenic pathway.

## Acknowledgement

The authors thank Dr. Yue Yu for implementation of the US methods in our lab. The authors also thank Virginia Tech Advanced Research Computing for computing time and resources. This work was supported by the National Institutes of Health (grant R35GM133754) and USDA-NIFA (project VA-160092).

## Supporting Information Available

The following items are available free of charge.

- SI.pdf: a PDF file containing 13 Figures including central structures, per-residue secondary structure analysis, dipole moment, and SASA analysis of second- and third-most occupied clusters, hydrogen bond analysis for Gln22 and Asn23 with backbone amide groups, the sidechain difference distance matrix, convergence of Δ_fold_, detail of the free energy minima and representative snapshots, and salt bridge persistence as a function of progress along the reaction coordinate. Four Tables are provided that include peptide-bond dipole moments, SASA, and sidechain dipole moments in the most-occupied cluster, and ΔG_fold_ of each variant over time.

## TOC Graphic

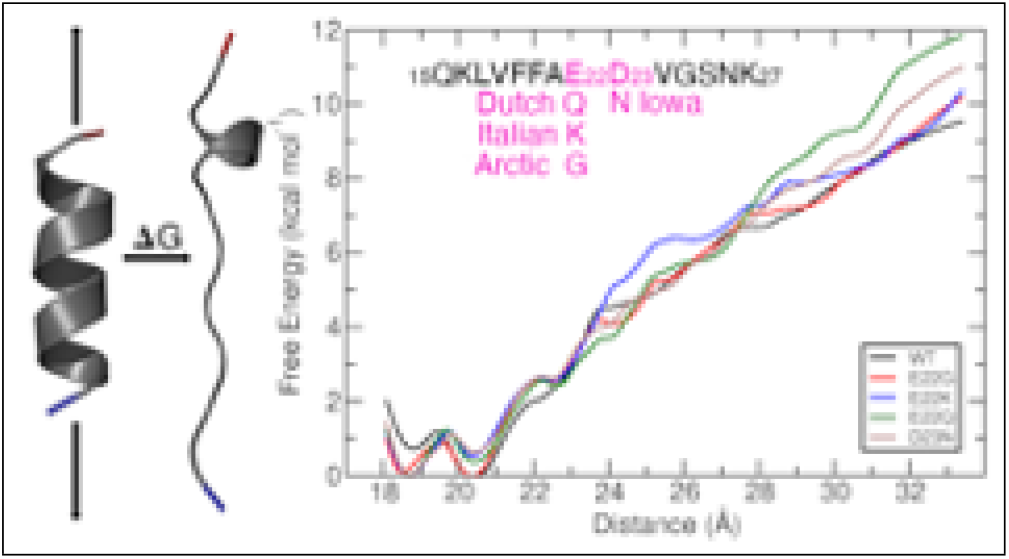

