## Supporting Information for "Effects of Familial Alzheimer’s Disease Mutations on the Folding Free Energy and Dipole-Dipole Interactions of the Amyloid *β*-Peptide"

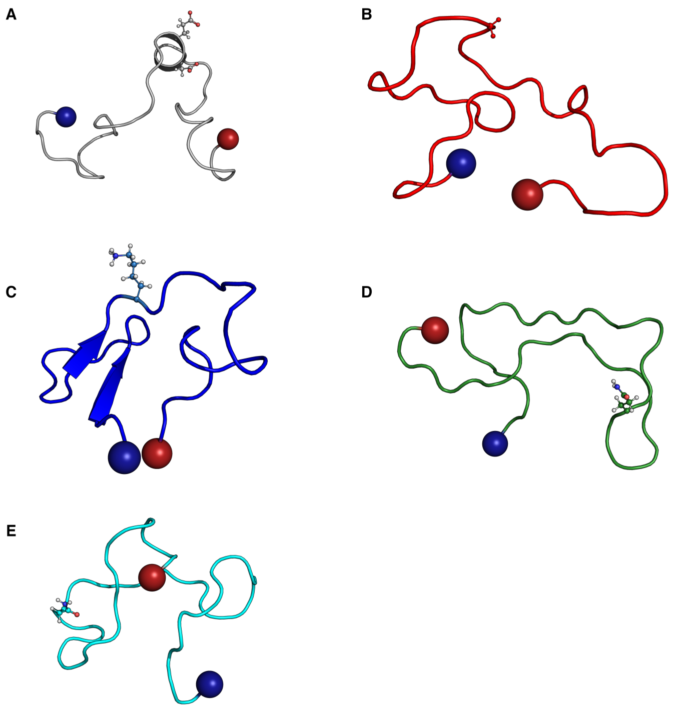


Figure S1: The central structure in each of the second most-occupied clusters for (A) WT, (B) E22G, (C) E22K, (D) E22Q, and (E) D23N peptides. Residues 22 and 23 are depicted in ball and stick. The N- and C-termini are colored blue and red, respectively.

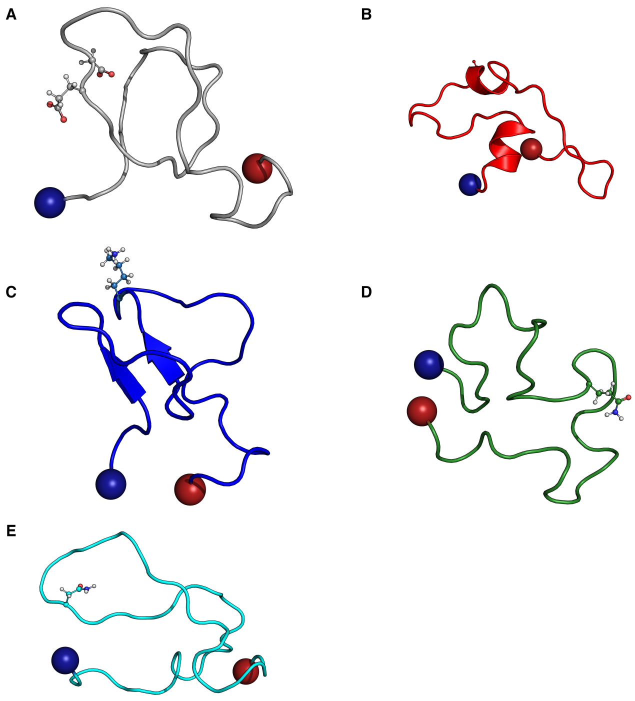


Figure S2: The central structure in each of the third most-occupied clusters for (A) WT, (B) E22G, (C) E22K, (D) E22Q, and (E) D23N peptides. Residues 22 and 23 are depicted in ball and stick. The N- and C-termini are colored blue and red, respectively.

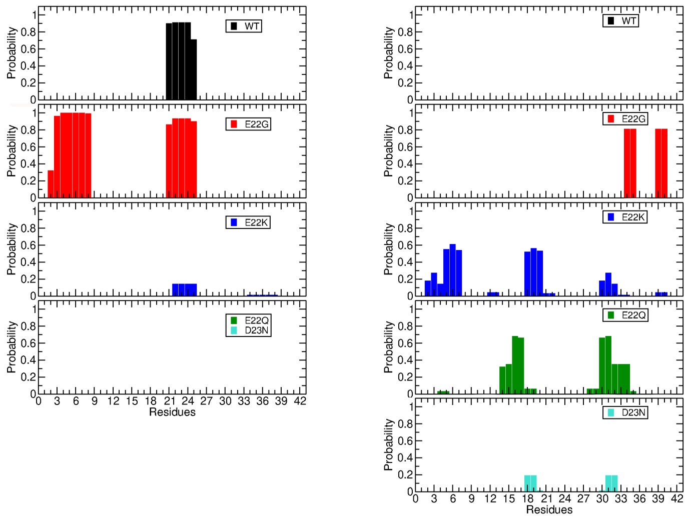


Figure S3: Secondary structure content in the second most-occupied cluster. On the left is the probability that the protein will form an $\alpha$-helix and the right shows $\beta$-sheet formation.

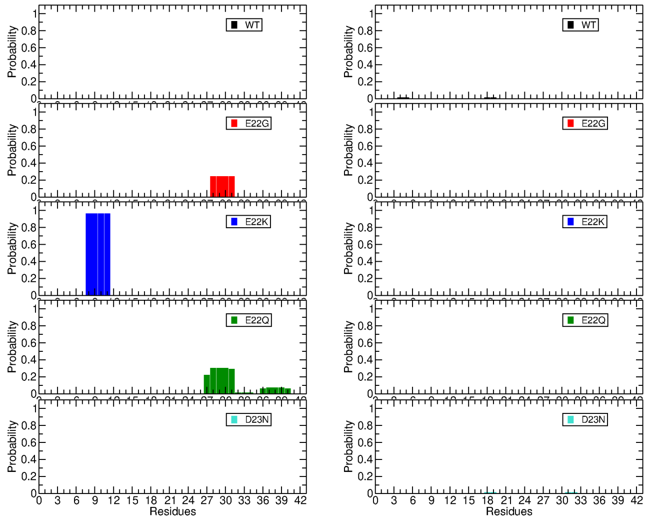

Figure S4: Secondary structure content in the third most-occupied cluster. On the left is the probability that the protein will form an $\alpha$-helix and the right shows $\beta$-sheet formation.

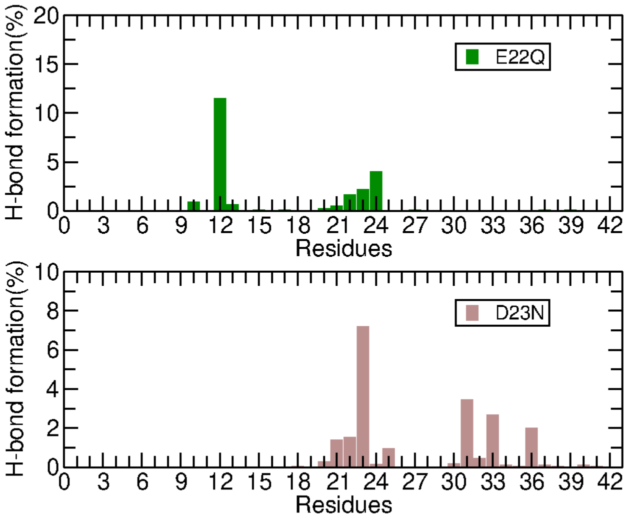


Figure S5: Frequency of hydrogen bond formation between the Gln22 (top) and Asn23 (bottom) sidechains with the backbone amide groups of each of the residues in the A$\beta_{42}$ peptide.

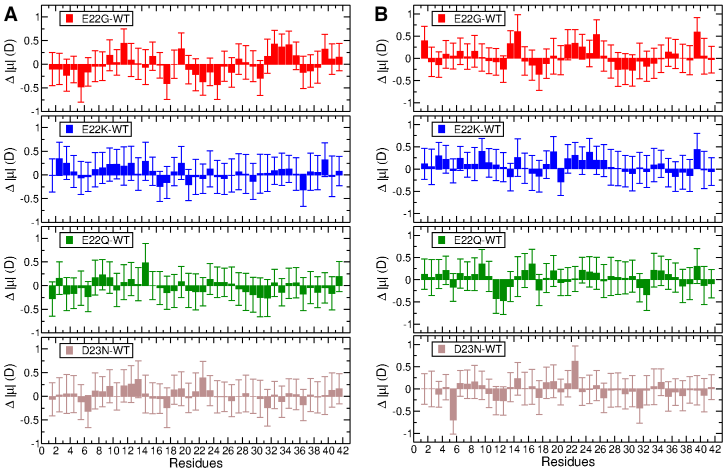


Figure S6: Difference in peptide-bond dipole moment as compared to the WT for (A) the second-most-occupied cluster and (B) the third-most-occupied cluster.

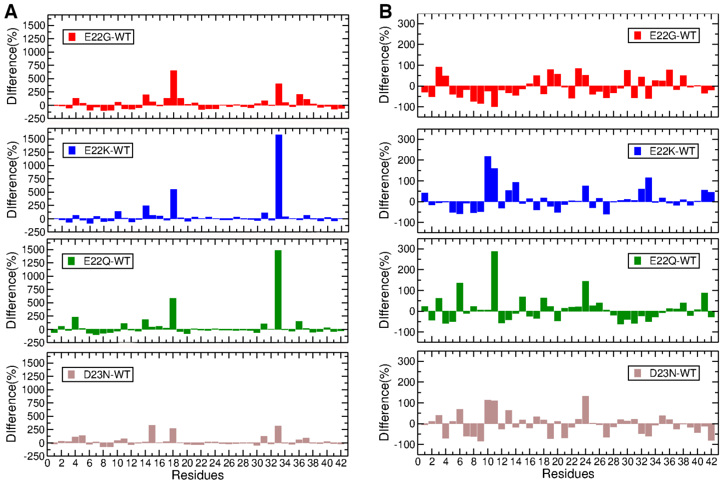


Figure S7: Percent difference in solvent-accessible surface area as compared to the WT in (A) the second-most-occupied cluster and (B) the third-most-occupied cluster.

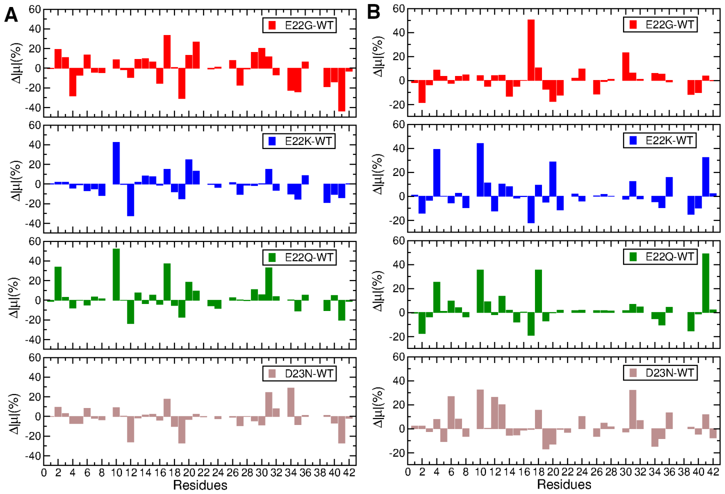


Figure S8: Percent difference in sidechain dipole moment as compared to the WT in (A) the second-most-occupied cluster and (B) the third-most-occupied cluster.

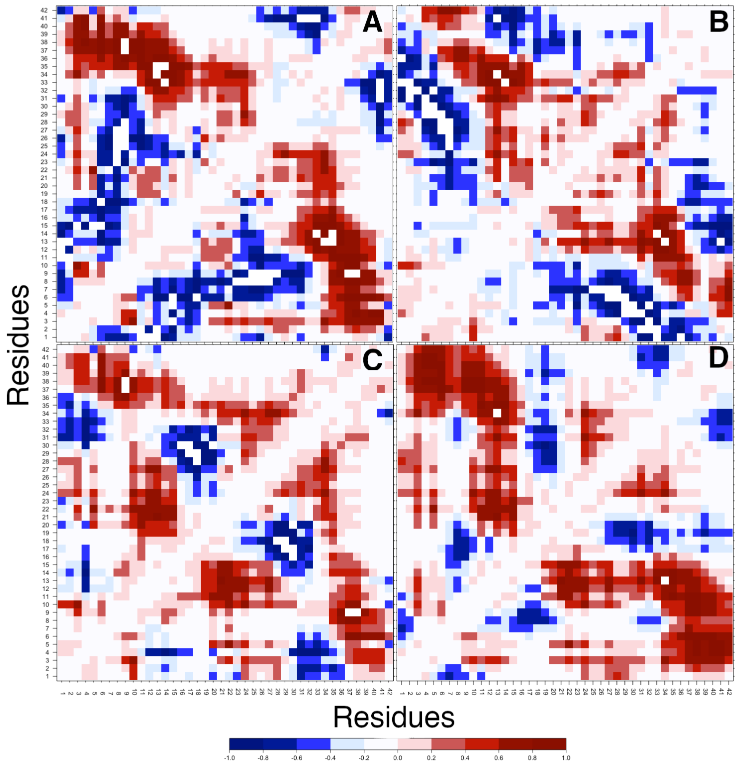


Figure S9: Sidechain distance matrix of the most-occupied cluster used to calculate tertiary structure interactions for the (A) WT, (B) E22G, (C) E22K, (D) E22Q, and (E) D23N peptides.

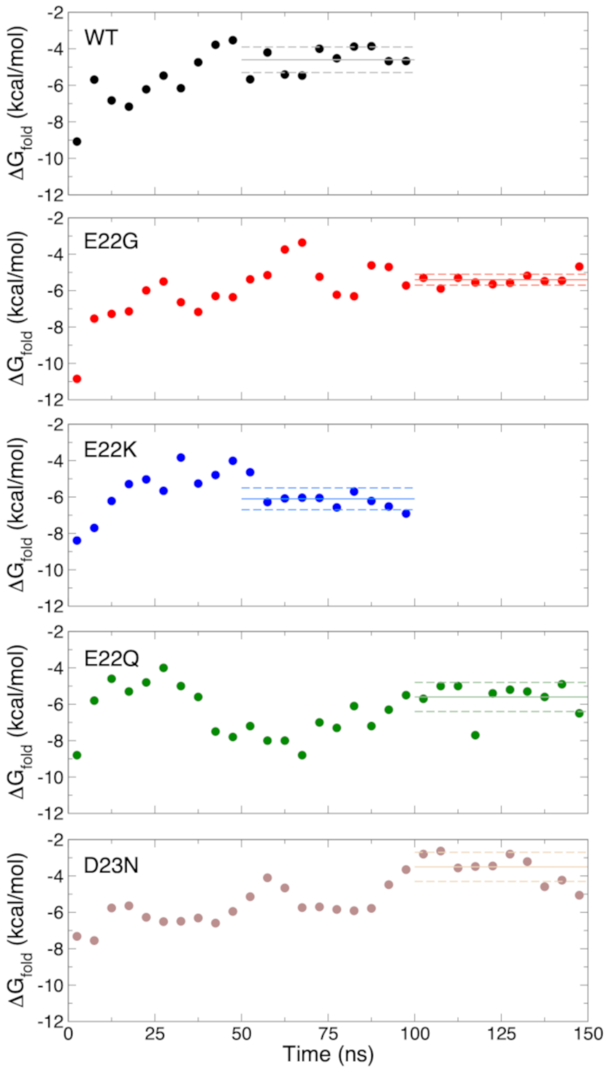


Figure S10: $\Delta G_{fold}$ over time, computed from 5-ns windows. Each data point is plotted in the middle of the corresponding time window. The solid, horizontal line in each panel correspond to the average value of $\Delta G_{fold}$ over the final 50 ns of simulation time, and the boundaries of the standard deviation are shown as dashed, horizontal lines.

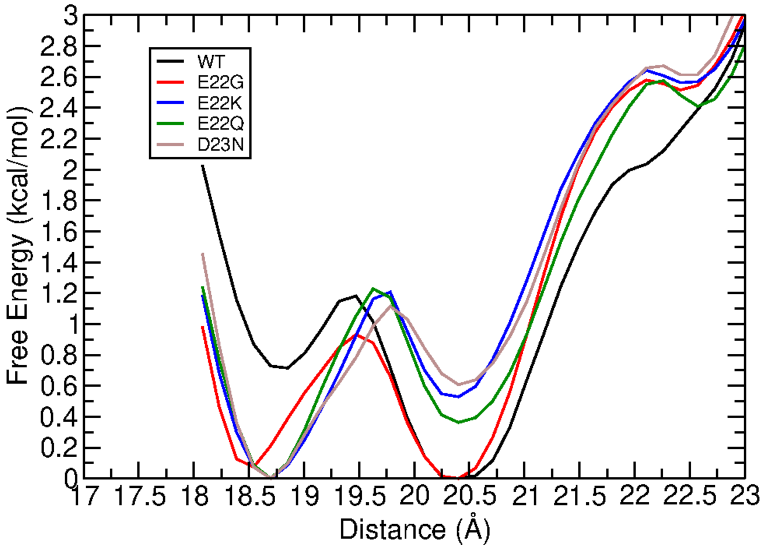


Figure S11: Zoomed in view of Figure 8 from main text that highlights the differences of the free energy surface between 18 and 21 Å along the reaction coordinate.

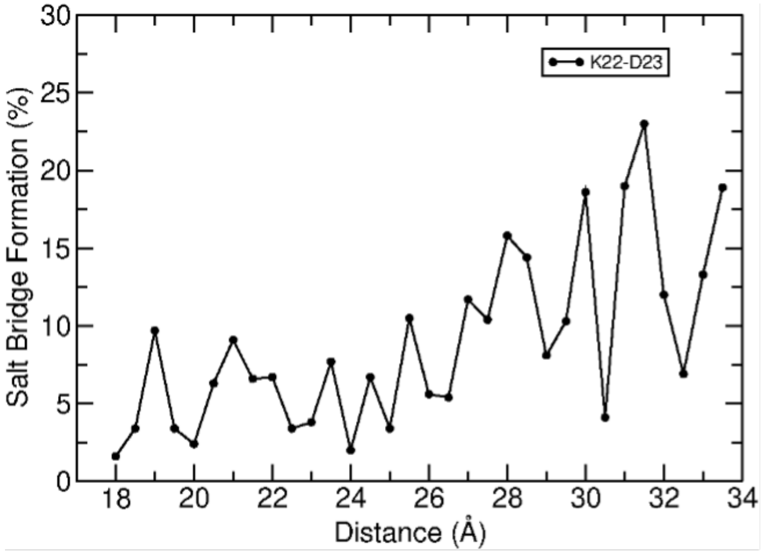


Figure S12: Percentage of time in which a salt bridge formed between Lys22 and Asp23 of E22K mutant at 298 K. Salt bridge occupancy was calculated using the minimum distance between the NZ atom of Lys22 and the OD1 or OD2 atom of Asp23. Salt bridges were considered formed when the distance between the atoms were less than or equal to 3.5 Å.

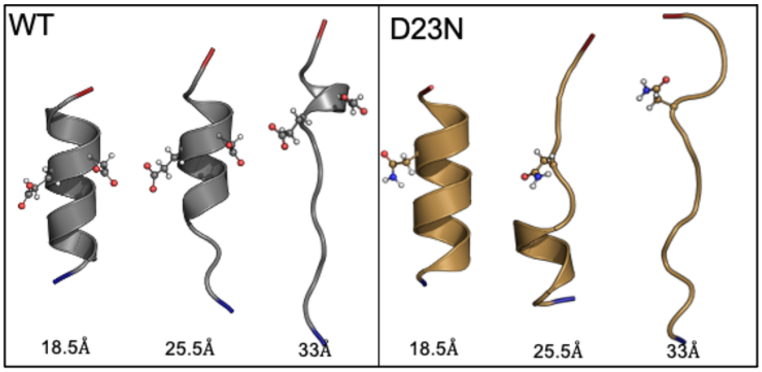


Figure S13: WT and D23N unfolding at the indicated reaction coordinate values during umbrella sampling simulations. E22 and D23 are highlighted in the WT as ball and stick with the N- and C-termini colored as blue and red, respectively. The N23 is drawn in ball and stick with the D23N.

Table S1: Full-length simulation peptide-bond dipole moments (D), computed from the snapshots in the most-occupied cluster. Two residues are shown for each entry to reflect the linkage between these two residues, which together constitute a net-neutral group for the calculation.

| Residues | WT | E22G | E22K | E22Q | D23N |
|---|---|---|---|---|---|
| 1-2 | 5.4 ± 0.2 | 5.7 ± 0.2 | 5.4 ± 0.2 | 5.4 ± 0.2 | 5.4 ± 0.2 |
| 2-3 | 4.9 ± 0.2 | 5.0 ± 0.2 | 5.3 ± 0.2 | 4.9 ± 0.2 | 5.0 ± 0.3 |
| 3-4 | 4.8 ± 0.2 | 4.9 ± 0.2 | 5.3 ± 0.2 | 4.8 ± 0.2 | 5.1 ± 0.3 |
| 4-5 | 5.0 ± 0.2 | 5.0 ± 0.2 | 5.5 ± 0.2 | 4.7 ± 0.2 | 5.0 ± 0.3 |
| 5-6 | 5.4 ± 0.2 | 5.6 ± 0.2 | 5.3 ± 0.2 | 5.3 ± 0.3 | 5.3 ± 0.3 |
| 6-7 | 5.2 ± 0.2 | 5.1 ± 0.2 | 5.1 ± 0.2 | 5.4 ± 0.2 | 5.0 ± 0.2 |
| 7-8 | 5.1 ± 0.3 | 5.0 ± 0.2 | 4.8 ± 0.3 | 5.0 ± 0.2 | 5.1 ± 0.2 |
| 8-9 | 5.1 ± 0.2 | 5.3 ± 0.2 | 5.0 ± 0.3 | 5.2 ± 0.2 | 5.1 ± 0.2 |
| 9-10 | 5.0 ± 0.3 | 4.8 ± 0.2 | 4.9 ± 0.2 | 4.9 ± 0.3 | 5.2 ± 0.3 |
| 10-11 | 4.9 ± 0.4 | 5.0 ± 0.2 | 4.9 ± 0.3 | 4.9 ± 0.2 | 4.9 ± 0.2 |
| 11-12 | 5.2 ± 0.2 | 5.1 ± 0.2 | 5.1 ± 0.3 | 5.3 ± 0.3 | 5.0 ± 0.3 |
| 12-13 | 5.1 ± 0.2 | 4.9 ± 0.2 | 5.0 ± 0.2 | 4.9 ± 0.2 | 5.1 ± 0.3 |
| 13-14 | 5.2 ± 0.2 | 5.5 ± 0.2 | 5.1 ± 0.3 | 5.0 ± 0.3 | 5.2 ± 0.4 |
| 14-15 | 5.5 ± 0.3 | 5.4 ± 0.2 | 5.1 ± 0.3 | 5.3 ± 0.3 | 5.0 ± 0.3 |
| 15-16 | 5.0 ± 0.2 | 4.9 ± 0.2 | 5.0 ± 0.2 | 5.0 ± 0.2 | 5.0 ± 0.3 |
| 16-17 | 4.8 ± 0.3 | 4.9 ± 0.2 | 5.0 ± 0.2 | 4.9 ± 0.2 | 5.1 ± 0.3 |
| 17-18 | 5.0 ± 0.2 | 4.9 ± 0.2 | 5.3 ± 0.2 | 5.2 ± 0.2 | 5.1 ± 0.3 |
| 18-19 | 5.0 ± 0.3 | 5.1 ± 0.2 | 5.2 ± 0.2 | 4.9 ± 0.3 | 5.3 ± 0.3 |
| 19-20 | 5.4 ± 0.3 | 5.1 ± 0.3 | 5.3 ± 0.3 | 5.1 ± 0.3 | 5.4 ± 0.3 |
| 20-21 | 5.3 ± 0.2 | 5.3 ± 0.3 | 5.4 ± 0.2 | 5.3 ± 0.2 | 5.3 ± 0.2 |
| 21-22 | 5.1 ± 0.3 | 5.2 ± 0.3 | 5.0 ± 0.2 | 5.2 ± 0.3 | 5.2 ± 0.3 |
| 22-23 | 5.0 ± 0.2 | 5.2 ± 0.2 | 5.3 ± 0.2 | 5.1 ± 0.2 | 5.6 ± 0.3 |
| 23-24 | 4.9 ± 0.2 | 5.0 ± 0.2 | 5.1 ± 0.2 | 4.9 ± 0.2 | 4.9 ± 0.3 |
| 24-25 | 5.0 ± 0.2 | 5.1 ± 0.2 | 5.3 ± 0.2 | 5.4 ± 0.2 | 5.2 ± 0.3 |
| 25-26 | 5.0 ± 0.3 | 5.5 ± 0.2 | 5.3 ± 0.3 | 5.3 ± 0.2 | 5.2 ± 0.2 |
| 26-27 | 5.1 ± 0.3 | 5.3 ± 0.2 | 5.5 ± 0.2 | 5.3 ± 0.3 | 5.3 ± 0.3 |
| 27-28 | 4.9 ± 0.2 | 4.9 ± 0.2 | 4.9 ± 0.2 | 4.9 ± 0.2 | 5.0 ± 0.3 |
| 28-29 | 5.2 ± 0.2 | 5.0 ± 0.2 | 5.2 ± 0.2 | 5.3 ± 0.3 | 5.2 ± 0.3 |
| 29-30 | 5.2 ± 0.3 | 5.0 ± 0.2 | 5.4 ± 0.2 | 5.4 ± 0.2 | 5.3 ± 0.3 |
| 30-31 | 5.1 ± 0.3 | 4.8 ± 0.2 | 5.2 ± 0.2 | 5.0 ± 0.2 | 5.0 ± 0.3 |
| 31-32 | 5.0 ± 0.2 | 5.0 ± 0.3 | 5.6 ± 0.2 | 4.8 ± 0.2 | 4.9 ± 0.3 |
| 32-33 | 5.1 ± 0.3 | 5.1 ± 0.2 | 5.2 ± 0.2 | 5.2 ± 0.2 | 5.2 ± 0.2 |
| 33-34 | 5.1 ± 0.2 | 5.0 ± 0.2 | 5.0 ± 0.2 | 5.2 ± 0.2 | 5.0 ± 0.3 |
| 34-35 | 5.3 ± 0.3 | 4.9 ± 0.2 | 4.9 ± 0.2 | 4.8 ± 0.3 | 4.9 ± 0.2 |
| 35-36 | 5.1 ± 0.3 | 5.1 ± 0.2 | 4.9 ± 0.2 | 5.0 ± 0.2 | 4.9 ± 0.2 |
| 36-37 | 5.2 ± 0.2 | 5.3 ± 0.2 | 5.3 ± 0.2 | 5.2 ± 0.2 | 5.2 ± 0.2 |
| 37-38 | 5.1 ± 0.3 | 5.4 ± 0.2 | 5.2 ± 0.2 | 5.2 ± 0.2 | 5.1 ± 0.2 |
| 38-39 | 4.9 ± 0.2 | 5.1 ± 0.2 | 5.0 ± 0.2 | 5.3 ± 0.3 | 5.1 ± 0.2 |
| 39-40 | 5.0 ± 0.3 | 5.4 ± 0.2 | 5.0 ± 0.3 | 5.1 ± 0.3 | 4.9 ± 0.3 |
| 40-41 | 5.0 ± 0.2 | 5.2 ± 0.3 | 5.1 ± 0.3 | 5.1 ± 0.3 | 5.3 ± 0.3 |
| 41-42 | 4.8 ± 0.3 | 5.1 ± 0.2 | 5.0 ± 0.2 | 5.1 ± 0.2 | 5.1 ± 0.2 |

Table S2: Solvent-accessible surface area (Å$^2$) of each residue in the full-length simulations, computed from the snapshots in the most-occupied cluster.

| Residue | WT | E22G | E22K | E22Q | D23N |
|---|---|---|---|---|---|
| Asp1 | 193.2 ± 32.3 | 100.8 ± 8.5 | 179.1 ± 11.9 | 149.9 ± 30.1 | 161.5 ± 46.3 |
| Ala2 | 77.0 ± 16.2 | 53.3 ± 6.3 | 65.2 ± 9.8 | 55.7 ± 25.5 | 87.7 ± 28.4 |
| Glu3 | 83.9 ± 30.1 | 179.8 ± 15.7 | 13.9 ± 7.1 | 81.5 ± 16.4 | 162.1 ± 52.1 |
| Phe4 | 157.3 ± 37.4 | 227.9 ± 10.0 | 127.4 ± 12.4 | 62.9 ± 36.2 | 160.9 ± 57.4 |
| Arg5 | 102.3 ± 27.0 | 67.0 ± 10.6 | 8.7 ± 11.6 | 109.4 ± 15.1 | 121.9 ± 36.0 |
| His6 | 89.5 ± 43.5 | 35.8 ± 8.6 | 87.1 ± 15.2 | 132.1 ± 16.6 | 139.2 ± 34.1 |
| Asp7 | 120.6 ± 31.1 | 65.0 ± 7.3 | 42.4 ± 16.3 | 108.2 ± 31.4 | 52.1 ± 26.9 |
| Ser8 | 100.9 ± 26.6 | 35.0 ± 13.3 | 87.6 ± 20.4 | 19.4 ± 8.1 | 39.3 ± 32.1 |
| Gly9 | 14.6 ± 20.5 | 14.9 ± 7.2 | 28.2 ± 23.1 | 63.5 ± 23.8 | 19.2 ± 12.6 |
| Tyr10 | 14.6 ± 20.5 | 14.9 ± 7.2 | 28.2 ± 23.1 | 63.5 ± 23.8 | 19.2 ± 12.6 |
| Glu11 | 43.5 ± 26.2 | 57.1 ± 12.6 | 50.7 ± 71.2 | 135.7 ± 39.6 | 82.2 ± 33.6 |
| Val12 | 59.6 ± 59.9 | 0.2 ± 2.5 | 89.1 ± 33.1 | 43.2 ± 26.5 | 128.5 ± 52.4 |
| His13 | 54.9 ± 50.8 | 110.5 ± 8.0 | 81.2 ± 23.0 | 97.3 ± 53.9 | 64.6 ± 31.3 |
| His14 | 97.3 ± 32.9 | 71.9 ± 7.3 | 134.5 ± 28.7 | 152.6 ± 47.1 | 164.9 ± 26.4 |
| Gln15 | 93.6 ± 22.6 | 54.8 ± 14.7 | 66.4 ± 35.2 | 48.3 ± 20.9 | 93.0 ± 38.3 |
| Lys16 | 62.0 ± 30.4 | 56.1 ± 10.0 | 74.3 ± 34.4 | 84.2 ± 30.2 | 153.1 ± 39.5 |
| Leu17 | 164.8 ± 45.4 | 193.6 ± 10.8 | 170.9 ± 20.1 | 175.9 ± 25.4 | 139.4 ± 31.2 |
| Val18 | 105.8 ± 33.5 | 151.2 ± 15.6 | 195.0 ± 10.1 | 125.5 ± 21.6 | 76.8 ± 47.3 |
| Phe19 | 166.6 ± 12.3 | 50.4 ± 10.9 | 71.8 ± 17.9 | 98.6 ± 23.2 | 96.0 ± 47.7 |
| Phe20 | 142.0 ± 24.2 | 217.7 ± 15.7 | 203.1 ± 16.5 | 76.0 ± 36.7 | 88.3 ± 36.6 |
| Ala21 | 134.1 ± 25.9 | 139.3 ± 33.4 | 71.0 ± 26.4 | 119.2 ± 36.8 | 97.9 ± 51.3 |
| Glu22 | 26.6 ± 8.7 | 72.8 ± 28.6 | 61.1 ± 13.6 | 48.0 ± 20.5 | 35.1 ± 22.3 |
| Asp23 | 93.4 ± 52.9 | 51.1 ± 17.6 | 131.7 ± 29.0 | 97.9 ± 43.1 | 120.0 ± 47.1 |
| Val24 | 102.1 ± 22.4 | 103.8 ± 16.5 | 38.9 ± 18.8 | 122.2 ± 26.4 | 82.1 ± 18.9 |
| Gly25 | 51.1 ± 9.6 | 31.1 ± 4.8 | 52.9 ± 11.1 | 56.7 ± 13.3 | 54.7 ± 16.7 |
| Ser26 | 81.4 ± 43.2 | 91.7 ± 14.6 | 143.0 ± 34.7 | 134.9 ± 22.6 | 160.2 ± 21.3 |
| Asn27 | 51.1 ± 9.6 | 31.1 ± 4.8 | 52.9 ± 11.1 | 56.7 ± 13.3 | 54.7 ± 16.7 |
| Lys28 | 85.0 ± 41.1 | 46.1 ± 9.1 | 96.5 ± 18.0 | 89.6 ± 29.7 | 87.4 ± 38.4 |
| Gly29 | 65.6 ± 15.8 | 64.5 ± 14.0 | 32.5 ± 12.8 | 24.3 ± 19.8 | 70.9 ± 19.6 |
| Ala30 | 60.2 ± 38.9 | 54.1 ± 13.6 | 34.5 ± 16.8 | 63.4 ± 31.3 | 66.6 ± 36.3 |
| Ile31 | 165.2 ± 34.1 | 134.4 ± 40.5 | 185.4 ± 22.1 | 144.7 ± 48.4 | 180.4 ± 26.7 |
| Ile32 | 65.6 ± 15.8 | 64.5 ± 14.0 | 32.5 ± 12.8 | 24.3 ± 19.8 | 70.9 ± 19.6 |
| Gly33 | 16.8 ± 12.5 | 11.0 ± 10.0 | 47.4 ± 9.3 | 34.9 ± 13.1 | 9.0 ± 13.6 |
| Leu34 | 83.7 ± 29.5 | 74.6 ± 26.1 | 46.5 ± 9.1 | 37.0 ± 24.1 | 51.6 ± 15.3 |
| Met35 | 95.5 ± 47.7 | 66.9 ± 21.8 | 135.5 ± 9.6 | 112.4 ± 35.7 | 116.2 ± 44.7 |
| Val36 | 177.9 ± 25.2 | 145.5 ± 20.5 | 34.9 ± 15.4 | 128.7 ± 24.0 | 101.2 ± 23.0 |
| Gly37 | 16.3 ± 14.4 | 51.7 ± 11.0 | 34.6 ± 15.3 | 29.7 ± 23.9 | 73.1 ± 15.8 |
| Gly38 | 15.9 ± 13.8 | 77.1 ± 13.5 | 6.8 ± 7.8 | 30.2 ± 19.1 | 48.0 ± 10.5 |
| Val39 | 16.8 ± 12.5 | 11.0 ± 10.0 | 47.4 ± 9.3 | 34.9 ± 13.1 | 9.0 ± 13.6 |
| Val40 | 55.5 ± 23.8 | 112.3 ± 27.0 | 142.6 ± 26.4 | 165.6 ± 34.5 | 71.2 ± 47.7 |
| Ile41 | 62.8 ± 35.0 | 129.2 ± 20.1 | 147.3 ± 35.2 | 100.3 ± 49.9 | 98.6 ± 41.0 |
| Ala42 | 124.0 ± 43.9 | 166.9 ± 11.9 | 63.2 ± 13.3 | 98.7 ± 36.9 | 85.6 ± 39.8 |

Table S3: Sidechain dipole moments (D) of each residue in the full-length simulations, computed from the snapshots in the most-occupied cluster. Glycine residues are omitted as they lack sidechain atoms.

| Residue | WT | E22G | E22K | E22Q | D23N |
|---|---|---|---|---|---|
| Asp1 | 5.1 ± 0.3 | 5.0 ± 0.3 | 5.1 ± 0.3 | 5.1 ± 0.3 | 5.1 ± 0.3 |
| Ala2 | 0.7 ± 0.1 | 0.7 ± 0.1 | 0.7 ± 0.1 | 0.7 ± 0.1 | 0.7 ± 0.1 |
| Glu3 | 6.7 ± 0.4 | 6.6 ± 0.5 | 6.8 ± 0.4 | 6.8 ± 0.4 | 6.6 ± 0.4 |
| Phe4 | 1.3 ± 0.4 | 0.8 ± 0.3 | 0.8 ± 0.3 | 1.5 ± 0.4 | 0.9 ± 0.4 |
| Arg5 | 6.0 ± 0.7 | 7.2 ± 0.5 | 6.5 ± 0.5 | 6.3 ± 0.6 | 6.4 ± 0.7 |
| His6 | 5.5 ± 0.5 | 5.1 ± 0.4 | 5.3 ± 0.5 | 5.3 ± 0.6 | 6.0 ± 0.6 |
| Asp7 | 5.2 ± 0.3 | 5.2 ± 0.3 | 5.2 ± 0.3 | 5.3 ± 0.3 | 5.3 ± 0.4 |
| Ser8 | 2.6 ± 0.3 | 2.5 ± 0.3 | 2.4 ± 0.4 | 2.1 ± 0.2 | 2.5 ± 0.3 |
| Tyr10 | 2.2 ± 0.6 | 2.0 ± 0.4 | 2.2 ± 0.6 | 2.4 ± 0.5 | 2.3 ± 0.5 |
| Glu11 | 6.7 ± 0.4 | 5.9 ± 0.3 | 6.7 ± 0.4 | 6.6 ± 0.4 | 6.7 ± 0.6 |
| Val12 | 0.6 ± 0.2 | 0.6 ± 0.2 | 0.5 ± 0.2 | 0.6 ± 0.2 | 0.6 ± 0.2 |
| His13 | 5.7 ± 0.6 | 5.4 ± 0.4 | 6.0 ± 0.6 | 6.1 ± 0.5 | 5.7 ± 0.7 |
| His14 | 5.3 ± 0.4 | 4.9 ± 0.4 | 5.4 ± 0.4 | 5.3 ± 0.4 | 5.5 ± 0.6 |
| Gln15 | 5.4 ± 0.5 | 5.6 ± 0.4 | 5.6 ± 0.6 | 5.5 ± 0.5 | 5.6 ± 0.6 |
| Lys16 | 10.4 ± 1.3 | 10.4 ± 1.2 | 10.4 ± 1.3 | 10.3 ± 1.2 | 10.3 ± 1.2 |
| Leu17 | 0.8 ± 0.3 | 1.1 ± 0.3 | 0.7 ± 0.3 | 0.8 ± 0.3 | 0.7 ± 0.3 |
| Val18 | 0.7 ± 0.2 | 0.6 ± 0.2 | 0.5 ± 0.2 | 0.6 ± 0.2 | 0.6 ± 0.2 |
| Phe19 | 0.8 ± 0.3 | 0.8 ± 0.3 | 0.8 ± 0.3 | 0.9 ± 0.3 | 0.8 ± 0.3 |
| Phe20 | 0.7 ± 0.3 | 0.8 ± 0.3 | 0.7 ± 0.3 | 0.7 ± 0.3 | 0.8 ± 0.3 |
| Ala21 | 0.7 ± 0.1 | 0.8 ± 0.2 | 0.7 ± 0.1 | 0.7 ± 0.1 | 0.7 ± 0.1 |
| Glu22 | 6.9 ± 0.4 | N/A | 10.0 ± 1.3 | 5.6 ± 0.5 | 6.5 ± 0.5 |
| Asp23 | 5.2 ± 0.4 | 5.3 ± 0.3 | 5.1 ± 0.3 | 5.2 ± 0.3 | 6.4 ± 0.4 |
| Val24 | 0.6 ± 0.2 | 0.7 ± 0.2 | 0.6 ± 0.2 | 0.6 ± 0.2 | 0.7 ± 0.2 |
| Ser26 | 2.4 ± 0.4 | 2.1 ± 0.3 | 2.3 ± 0.3 | 2.3 ± 0.3 | 2.2 ± 0.3 |
| Asn27 | 5.9 ± 0.5 | 5.9 ± 0.4 | 6.4 ± 0.4 | 6.6 ± 0.5 | 6.2 ± 0.4 |
| Lys28 | 10.5 ± 1.2 | 10.5 ± 1.2 | 10.4 ± 1.2 | 10.5 ± 1.2 | 10.4 ± 1.2 |
| Ala30 | 0.8 ± 0.1 | 0.8 ± 0.1 | 0.7 ± 0.1 | 0.7 ± 0.1 | 0.7 ± 0.1 |
| Ile31 | 0.7 ± 0.2 | 0.7 ± 0.2 | 0.6 ± 0.2 | 0.8 ± 0.2 | 0.7 ± 0.3 |
| Ile32 | 0.6 ± 0.2 | 0.6 ± 0.2 | 0.6 ± 0.2 | 0.7 ± 0.2 | 0.6 ± 0.2 |
| Leu34 | 0.8 ± 0.3 | 0.8 ± 0.3 | 0.7 ± 0.3 | 0.7 ± 0.3 | 0.9 ± 0.4 |
| Met35 | 2.5 ± 0.4 | 3.1 ± 0.4 | 2.7 ± 0.5 | 2.6 ± 0.5 | 2.9 ± 0.5 |
| Val36 | 0.6 ± 0.2 | 0.6 ± 0.2 | 0.5 ± 0.2 | 0.6 ± 0.2 | 0.6 ± 0.2 |
| Val39 | 0.7 ± 0.2 | 0.6 ± 0.2 | 0.7 ± 0.2 | 0.6 ± 0.2 | 0.7 ± 0.2 |
| Val40 | 0.7 ± 0.2 | 0.6 ± 0.2 | 0.6 ± 0.2 | 0.6 ± 0.2 | 0.6 ± 0.2 |
| Ile41 | 0.6 ± 0.2 | 0.5 ± 0.2 | 0.7 ± 0.2 | 0.6 ± 0.2 | 0.6 ± 0.2 |
| Ala42 | 1.0 ± 0.1 | 1.0 ± 0.1 | 1.0 ± 0.1 | 1.0 ± 0.4 | 1.0 ± 0.1 |

Table S4: $\Delta G_{fold}$ (kcal/mol) of the WT and mutants in each of the indicated time intervals.

| Time Interval (ns) | WT | E22Q | E22K | E22G | D23N |
|---|---|---|---|---|---|
| 0-5 | -9.08 | -8.8 | -8.39 | -10.85 | -7.32 |
| 5-10 | -5.69 | -5.8 | -7.70 | -7.54 | -7.55 |
| 10-15 | -6.83 | -4.6 | -6.22 | -7.28 | -5.76 |
| 15-20 | -7.17 | -5.3 | -5.29 | -7.14 | -5.64 |
| 20-25 | -6.22 | -4.8 | -5.03 | -5.99 | -6.27 |
| 25-30 | -5.47 | -4.0 | -5.66 | -5.50 | -6.51 |
| 30-35 | -6.16 | -5.0 | -3.83 | -6.64 | -6.49 |
| 35-40 | -4.74 | -5.6 | -5.26 | -7.17 | -6.31 |
| 40-45 | -3.78 | -7.5 | -4.79 | -6.3 | -6.59 |
| 45-50 | -3.53 | -7.8 | -4.01 | -6.36 | -5.95 |
| 50-55 | -5.67 | -7.2 | -4.64 | -5.38 | -5.14 |
| 55-60 | -4.20 | -8.0 | -6.28 | -5.15 | -4.10 |
| 60-65 | -5.41 | -8.0 | -6.08 | -3.74 | -4.66 |
| 65-70 | -5.47 | -8.8 | -6.04 | -3.36 | -5.74 |
| 70-75 | -4.00 | -7.0 | -6.05 | -5.24 | -5.70 |
| 75-80 | -4.52 | -7.3 | -6.57 | -6.23 | -5.84 |
| 80-85 | -3.88 | -6.1 | -5.70 | -6.31 | -5.91 |
| 85-90 | -3.86 | -7.2 | -6.22 | -4.61 | -5.78 |
| 90-95 | -4.68 | -6.3 | -6.52 | -4.7 | -4.48 |
| 95-100 | -4.67 | -5.5 | -6.91 | -5.72 | -3.65 |
| 100-105 | | -5.7 | | -5.30 | -2.80 |
| 105-110 | | -5.0 | | -5.89 | -2.63 |
| 110-115 | | -5.0 | | -5.3 | -3.56 |
| 115-120 | | -7.7 | | -5.55 | -3.48 |
| 120-125 | | -5.4 | | -5.65 | -3.45 |
| 125-130 | | -5.2 | | -5.57 | -2.80 |
| 130-135 | | -5.3 | | -5.18 | -3.21 |
| 135-140 | | -5.6 | | -5.48 | -4.59 |
| 140-145 | | -4.9 | | -5.45 | -4.23 |
| 140-150 | | -6.5 | | -4.68 | -5.06 |
